# Tryptophan metabolism reprogramming contributes to the prothrombotic milieu in mice and humans infected with SARS-CoV-2

**DOI:** 10.1101/2025.01.17.633602

**Authors:** Saravanan Subramaniam, Marc Arthur Napoleon, Saran Lotfollahzadeh, Mohamed Hassan Kamal, Helena Kurniawan, Murad Elsadawi, Devin Kenney, Florian Douam, Markus Bosmann, Stephen Whelan, Howard Cabral, Eric J Burks, Grace Zhao, Vijay Kolachalama, Katya Ravid, Vipul C. Chitalia

**Affiliations:** Department of Pharmacology and Toxicology, School of Pharmacy, Massachusetts College of Pharmacy and Health Sciences, Boston, MA 02115, USA; Pulmonary Center, Department of Medicine, Chobanian & Avedisian School of Medicine, Boston University, Boston, MA 02118, USA; Renal Section, Department of Medicine, Chobanian & Avedisian School of Medicine, Boston University, Boston, MA 02118, USA; Department of Pathology and Laboratory Medicine, Chobanian & Avedisian School of Medicine, Boston University, Boston, MA 02118, USA; Department of Virology, Immunology and Microbiology, Chobanian & Avedisian School of Medicine, Boston University, Boston, MA 02118, USA; National Emerging Infectious Diseases Laboratories, Boston University, Boston, MA 02118, USA; Center for Thrombosis and Hemostasis, University Medical Center of the Johannes Gutenberg-University, Mainz, Germany; Chemical Instrumentation Center, Department of Chemistry, Boston University, Boston, MA, 02118, USA; Department of Biostatistics, School of Public Health, Boston University, MA, 02118, USA; Computational Medicine, Chobanian & Avedisian School of Medicine, Boston University, Boston, MA 02118, USA; Department of Medicine and Whitaker Cardiovascular Institute, Chobanian & Avedisian School of Medicine, Boston University, Boston, MA 02118, USA; Veterans Affairs Boston Healthcare System, Boston MA 02118, USA

**Author notes:** Shared first authors (Saravanan Subramaniam, Marc Arthur Napoleon). Corresponding author: Vipul Chitalia, M.D., Ph.D., Director, Center of Cross-Organ Vascular Pathology, Department of Medicine, Boston University Medical Center, X-545 Boston, MA 02118, USA, (P) (F).

**Keywords:** COVID-19, humanized ACE2 mice, tryptophan catabolism, kynurenine pathway

## Abstract

SARS-CoV-2 infection disturbs the coagulation balance in the blood, triggering thrombosis and contributing to organ failure. The role of prothrombotic metabolites in COVID-19-associated coagulopathy remains elusive. Leveraging K18-hACE2 mice infected with SARS-CoV-2, we observed higher levels of the tryptophan metabolite, kynurenine, compared to controls. SARS-CoV-2 infected mice showed a significant upregulation of enzymes controlling Kynurenine biogenesis, such as indoleamine 2,3-dioxygenase (IDO-1) and tryptophan 2,3-dioxygenase levels in kidneys and liver, respectively, as well as changes in the enzymes involved in kynurenine catabolism, including kynurenine monooxygenase and kynurinase. Consistent with the agonistic role of these metabolites in Aryl Hydrocarbon Receptor (AHR) signaling, AHR activation and its downstream mediator, tissue factor (TF), a highly potent procoagulant factor, was observed in endothelial cells (ECs) of lungs and kidneys of infected mice. These findings were validated in humans, where compared to controls, sera of COVID-19 patients showed increased levels of Kynurenine, kynurenic acid, anthranilic acid, and quinolinic acid. Activation of the AHR-TF axis was noted in the kidneys and lungs of COVID-19 patients, and COVID-19 sera showed higher IDO-1 activity than controls. Levels of Kyn in COVID-19 patients correlated strongly with the TF-inducing activity of COVID-19 sera on ECs. A specific IDO-1 inhibitor or AHR inhibitor separately or in combination suppressed COVID-19 sera-induced TF activity in ECs. Together, we identified IDO-1 as upregulated by SARS-CoV-2 infection, resulting in augmented Kyn and its prothrombotic catabolites, thereby suggesting the Kyn-AHR-TF axis as possibly a new diagnostic and/or therapeutic target.

**Key points:**

1. SARS-CoV-2-infection upregulates kynurenine biogenesis in the liver and diminishes kynurenine catabolism in the lungs and kidneys.
2. An increase in kynurenine stimulates the AHR-TF axis in the microvasculature in COVID-19 patients, which is inhibited by pharmacological manipulation.

## Introduction

SARS-CoV-2 infection induces a profoundly prothrombotic milieu by creating an imbalance between pro- and anti-coagulant factors in the blood. This perturbation drives several thrombotic complications in the arterial and venous beds of these patients, resulting in potentially fatal complications such as pulmonary embolism, deep vein thrombosis, stroke, and coronary artery syndrome ^1–3^. In addition to macrovascular thrombosis, many studies demonstrated microthrombi in the alveolar and renal capillary beds in mouse models ^4,5^ and humans ^6,7^. Although thrombotic complications are common in other viral infections, SARS-CoV-2 infection differentiates itself due to a higher risk of thromboembolic phenomenon. COVID-19 results in approximately 9-fold higher occurrence of alveolar microthrombi as compared to H1N1 patients ^8^ ^9,10^. Because microthrombi are an important contributor to renal and respiratory failure, it is imperative to identify mediators of microvascular thrombosis in these patients.

COVID-19-associated coagulopathy involves complex interactions between innate immunity, extrinsic coagulation and fibrinolytic pathways, and the vascular endothelium, platelets, and white blood cells (WBCs) ^1,11,12^. Tissue factor (TF) is the primary trigger of the extrinsic coagulation cascade during infection ^1^. Blood from COVID-19 patients shows extrinsic coagulation cascade activation, as evidenced by abnormal prothrombin time, and activated partial thromboplastin time prolongation ^2,13,14^, supporting TF activation. TF is a membrane protein, and its activation on the cell surface in different cell types, including endothelial cells (ECs), triggers an extrinsic coagulation cascade ^10,15^.

A growing body of research points to the involvement of ‘*thrombolome*’—specific metabolites with prothrombotic properties—in thrombosis in other disease models. Previous studies from our lab and that of others showed that L-tryptophan (Trp)’s metabolites, such as L-Kynurenine (Kyn) and its other catabolites, augment thrombosis in animal models and patients with chronic kidney disease (CKD) ^16–20^ and in some cancer models ^21^. A large portion of dietary Trp (>80%) is converted in the liver to Kyn by a rate-limiting step catalyzed by tryptophan 2, 3-dioxygenase (TDO) and indoleamine 2,3-dioxygenase (IDO). Kyn is further degraded by Kynureninase or L-Kynurenine hydrolase (Kyn), Kynurenine aminotransferases (KATs) and 3-hydroxyanthranilate oxidase (3-HAO) to different metabolites such as anthranilic acid (AA), kynurenic acid (KYNA), xanthurenic acid (XA), quinolinic acid (QA) and picolinic acid (PA) to generate nicotinamide adenine dinucleotide (NAD+) required for cellular metabolism ^22^ (**Figure 1A**). The enzymes regulating Kyn biogenesis and degradation are expressed in different organs and cell types. ^23^. TDO is mainly expressed in the liver ^24^. IDO expression is limited to dendritic cells ^25^, vascular smooth muscle (vSMC) ^26^, and ECs ^27^.

**Figure 1:**
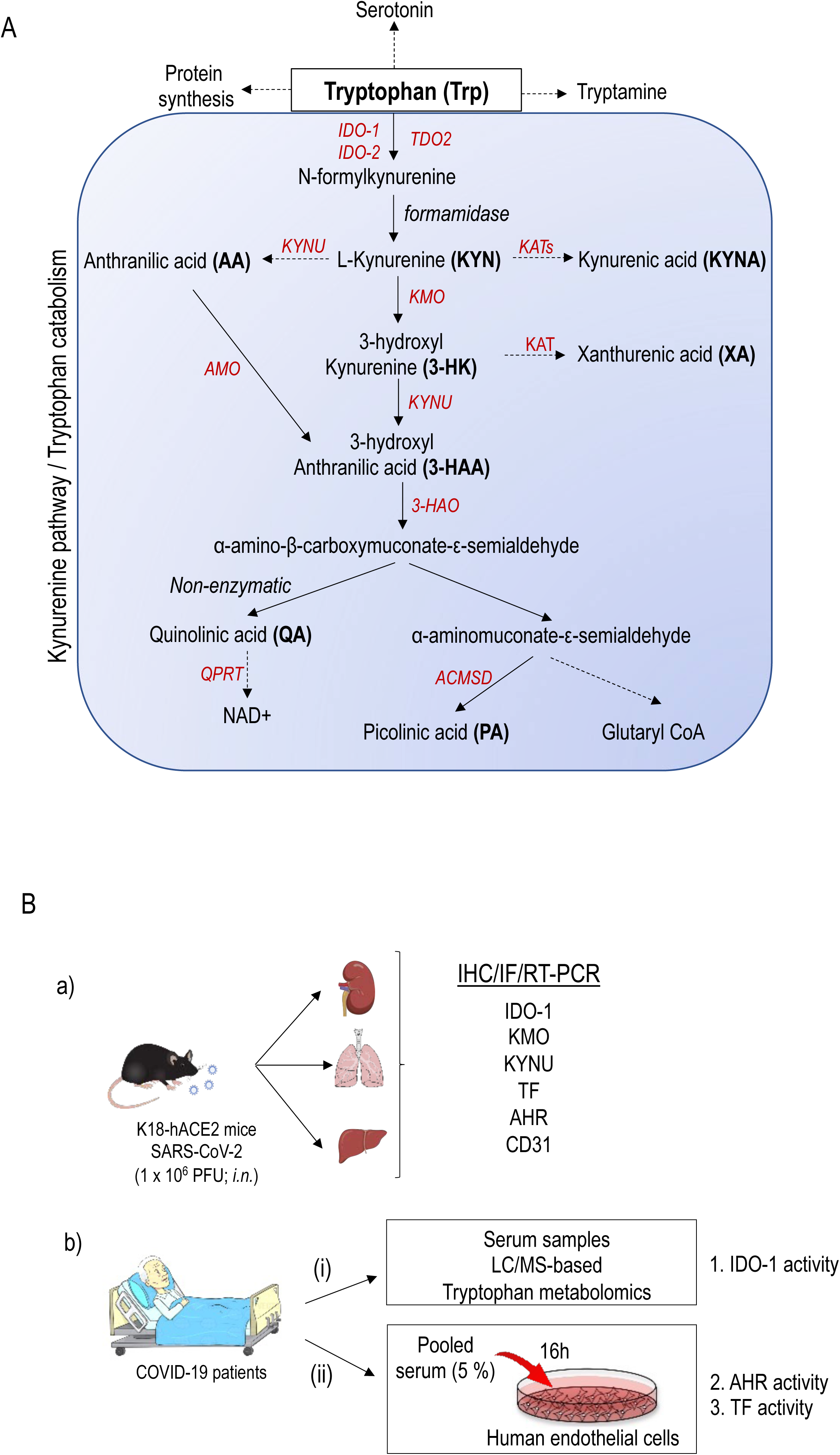
Schematic overview of Tryptophan catabolism. **(A)** Diet with tryptophan is degraded into two major segments. Ninety percent is absorbed and is converted to the N-Formylkynurenine in the liver via two important enzymes, IDO-1/IDO-2 and TDO2. The remaining 10 % is converted into indol. N-Formylkynurenine is converted to KYN via (Aryl) formamidase. However, N-formylkynurenine is converted to kynurenic acid via (KAT-I/-II/-III). KYN is converted to 3-HK via KMO. 3-HK is further degraded to XA, and to 3-HAA. The latter is catalyzed via Kynureninase (KYNU). AA, anthranilic acid; ACMSD, aminocarboxymuconate-semialdehyde-decarboxylase; AMO, anthranilate 3-monooxygenase; 3-HAA, 3-hydroxyanthranilic acid; 3-HAO, 3-hydroxyanthranilate 3,4-dioxygenase; 3-HK, 3-hydroxykynurenine; IDO-1 and or, IDO-2, indoleamine 2,3-dioxygenase-1 and −2; KATs, kynurenine aminotransferase enzymes; KMO, kynurenine monooxygenase; KYN, kynurenine; KYNA, kynurenic acid; NAM, nicotinamide adenine mononucleotide; PA, picolinic acid; QA, quinolinic acid; QPRT, quinolinate phosphoribosyltransferase; TDO, tryptophan-2,3-dioxygenase; Trp, Tryptophan; XA, xanthurenic acid. Enzymatic reactions are shown in *italics.* **(B) Schematic presentation of the experimental design**. **(a)** Humanized transgenic ACE2 expressing mice were inoculated intranasally with SARS-COV-2 and at specified time period, the mice were sacrificed, and organs (kidney, lung, and liver) and blood samples were harvested. Samples were subjected to targeted tryptophan metabolites. **(b)** Serum samples from COVID-19 patients were examined for **(i)** tryptophan metabolites by LC-MS-based metabolomics and **(ii)** pooled serum (5%) exposure of human ECs, followed by AHR and TF activity assays.

Thomas et al. showed that the levels of Kyn were significantly higher in COVID-19 patients ^28^, suggesting a potential alteration in Trp metabolism. Kyn is a ligand for aryl hydrocarbon receptor (AHR). Kyn activates AHR signaling to upregulate TF in ECs and vSMC in other disease models ^18,21^. In the current study, we set out to examine the hypothesis that SARS-CoV-2 infection upregulates enzymes controlling Kyn biogenesis and/or possibly suppresses enzymes involved in Kyn catabolism in organs at risk of microvascular thrombosis. Leveraging genetically humanized mice infected with SARS-CoV-2 and samples of infected humans (**Figure 1B**), we examined lungs, liver, and kidneys and employed pharmacological manipulations to probe the targetability of the Kyn-AHR-TF axis.

## Methods

### Mice

Heterozygous transgenic humanized ACE2 (K18-hACE2) mice (strain: 2B6.Cg-Tg(K18-ACE2)2Prlmn/J) were obtained from the Jackson Laboratory (Bar Harbor, ME). Animals were maintained in Tecniplast green line individually ventilated cages (Tecniplast, Buguggiate, Italy). Mice were maintained on a 12:12 light cycle at 30-70% humidity and provided *ad libitum* water and standard chow diets (LabDiet, St. Louis, MO). The Boston University Biomedical Research approved all experimental procedures with animals, Institutional Biosafety Committee and Institutional Animal Care and Use Committee.

### SARS-CoV-2 propagation

SARS-CoV-2 propagation was carried out following previously established methods^29,30^. The SARS-CoV-2 isolate 2019-nCoV/USA-WA1/2020 (NCBI accession number: MN985325) was sourced from the Centers for Disease Control and Prevention (Atlanta, GA) and BEI Resources (Manassas, VA). 1×10^7^ African green monkey kidney Vero E6 cells (ATCC® CRL-1586™, American Type Culture Collection, Manassas, VA) were seeded in a T-175 flask one day prior to virus generation. The following day, the cells were infected with the virus diluted in 10 mL of Opti-MEM (ThermoFisher Scientific, Waltham, MA, USA) and incubated for 1 hour at 37°C to facilitate virus adsorption. Subsequently, 15 mL of DMEM containing 10% FBS and 1% penicillin/streptomycin was added to the cells and incubated overnight. The next day, the media was aspirated, cells were washed with 1X PBS, pH 7.5 (ThermoFisher Scientific), and 25 mL of fresh DMEM containing 2% FBS was added. The cells were then monitored for cytopathic effect (CPE), the media was harvested, filtered through a 0.22 μm filter, and concentrated using a sucrose gradient. The concentrated virus was suspended in sterile 1X PBS, pH 7.5, aliquoted, and stored at –80°C.

### SARS-CoV-2 titration via plaque assay

The titer of our viral stocks was assessed using a plaque assay method. Vero E6 cells were plated in a 12-well plate at a density of 2×10^5 cells per well. The following day, the cells were exposed to 10-fold serial dilutions of the viral stocks and allowed to incubate for 1 hour at 37°C to enable viral attachment. Subsequently, 1 mL of overlay media (composed of 1.2% Avicel (DuPont, Wilmington, DE, USA; RC-581) in DMEM supplemented with 2% FBS and 1% Pen/Strep) was added to each well. Three days later, the overlay media was removed, and the cells were fixed with 10% neutral buffered formalin (ThermoFisher Scientific) for 1 hour at room temperature. After fixation, the formalin was removed, and the cells were stained with 0.1% crystal violet (Sigma-Aldrich) in 10% ethanol/water for 30 minutes at room temperature. Following staining, the crystal violet was removed, cells were washed with water, and plaque-forming units were enumerated to determine viral titers.

### SARS-CoV-2 inoculation of mice

K18-hACE2 transgenic mice of both sexes (aged 12-16 weeks) were intranasally administered with 1×10^6 plaque-forming units (PFU) of SARS-CoV-2 in 50 μL of sterile 1X PBS, or sham-inoculated. The inoculations were conducted under 1-3% isoflurane anesthesia. Mice were euthanized at predetermined time points for sample collection (2dpi, 4dpi, and 7dpi), or earlier if they met euthanasia criteria (as defined by an IACUC-approved clinical scoring system).

### Cell culture

Human umbilical vein endothelial cells (HUVEC)-TERT2 and HUVECs were cultured as described before ^18^. In brief, HUVEC-TERT2 cells were cultured in RPMI medium1640 (Cat. No. 11875-093, GIBCO, MA), containing 10 % fetal bovine serum, 1% penicillin/streptomycin in 5% CO2 at 37°C. HUVECs (Cat. no. C2517A; Lonza, Walkersville, MD) were cultured in EBM^TM^-2 Basal Medium (CC-3156) and EGMTM-2 SingleQuots^TM^ Supplements (CC-4176) with 1% penicillin/streptomycin in 5% CO2 at 37°C.

### COVID-19 sera and autopsy samples

Sera samples and kidney autopsies from COVID-19 patients were received from Boston Medical Center Biorepository (IRB: H26367 and IBC: 20-1459). Samples were collected from inpatient and ambulatory patients who provided informed consent to participate in the study. Participants were all confirmed positive for SARS-CoV-2 infection and were either admitted to the hospital or seen in an ambulatory setting for treatment.

To this end, we recruited 29 patients at Boston Medical Center (BMC) with confirmed COVID-19 of nasal epithelial PCR; most patients were male and had a mean age of 59 years. These were mainly non-Hispanic white people who had a variety of comorbid conditions, such as diabetes, hypertension, coronary artery disease, and stroke. Control patients admitted to BMC over the same time were matched with members of this group. Most control patients were females of black ethnicity and had comparable comorbidities; there were no statistical differences in age between the control and COVID-19 groups (see Table 1).

**Table 1.** Baseline characteristics of patients.

|  | Control (N= 20) | COVID-19 (N= 20) | Significance (P value) |
| --- | --- | --- | --- |
| Age Mean (SD) | 55.3±16.5 | 61.4±18.7 | 0.281 |
| Sex number (%) |  |  |  |
| Male | 13 (65%) | 15 (75%) |  |
| Female | 7 (35%) | 5 (25%) |  |
| Ethnicity |  |  |  |
| Hispanic (%) | 5 | 20 |  |
| Non-Hispanic (%) | 45 | 30 |  |
| Race |  |  |  |
| Caucasians (%) | 17.5 | 20 |  |
| African Americans (%) | 25 | 10 |  |
| Latino (%) | 5 | 15 |  |
| Asian (%) | - | 2.5 |  |
| Others# (%) | 2.5 | 2.5 |  |
| Comorbidities |  |  |  |
| HTN (%) | 40 (8/20) | 50 (10/20) | 0.525 |
| DM (%) | 20 (4/20) | 20 (4/20) | 1.00 |
| CAD (%) | 20 (4/20) | 5 (1/20) | 0.341 |
| CHF (%) | 30 (6/20) | 10 (2/20) | 0.235 |
| CKD (%) | 100 (20/20) | 100 (20/20) | 1.00 |
| Asthma (%) | 25 (5/20) | 20 (4/20) | 1.00 |
| COPD (%) | 10 (2/20) | 15 (3/20) | 1.00 |
| Cirrhosis (%) | 0 (0/20) | 10 (2/20) | 0.487 |
| Stroke (%) | 15 (3/20) | 15 (3/20) | 1.00 |
| Baseline vitals |  |  |  |
| Pulse | 94.4±21.7 | 88.8±12.1 | 0.326 |
| SBP | 135.9±22.3 | 130.4±21.9 | 0.432 |
| DBP | 78.8±15.3 | 77.8±8.7 | 0.889 |
| SaO2 | 92.1±21.6 | 57.9±47.8 | 0.007 |
| Temperature | 98.7±1.3 | 98.6±1.1 | 0.709 |
| Baseline labs |  |  |  |
| Hb | 10.8±3.2 | 12.1±2.8 | 0.173 |
| WBC | 9.7±5.7 | 8.4±3.9 | 0.413 |
| Platelets | 281.2±161.7 | 220.5±103.8 | 0.165 |
| BUN (gm/dL) | 16.7±9.8 | 17.1±9.9 | 0.892 |
| Creatine (mg/dL) | 1.0±0.4 | 0.98±0.43 | 0.909 |
| Na (mEq/L) | 138.3±2.5 | 137.6±4.9 | 0.577 |
| K (mEq/L) | 4.2±0.7 | 4.3±0.8 | .833 |
| CL (mEq/L) | 105.4±4.3 | 104.5±5.1 | .531 |
| HCO<sub>3</sub> (mEq/L) | 22.3±3.9 | 22.5±3.5 | .866 |
| D-dimer | 1810 ±617 | 4870.4 ±1057 | .0182 |
| Fibrinogen | 382 ±218 | 412.9 ±103 | .716 |

### LC/MS-based metabolomics study

Sera samples from COVID-19 patients and COVID-19-negative patients were used for LC/MS-based metabolomics study as described before ^31^. In brief, 200 µL of serum samples were subjected to solid phase extraction using methanol and 1% formic acid. The analytes were eluted with 1% ammonium peroxide in methanol/water (50:50). The elutes were dried under nitrogen and reconstitute (150 µL) with 5 mmol/l ammonium acetate solution. For standards, ten metabolites were weighed, dissolved in methanol, and diluted to specified concentrations in 5% bovine serum albumin/phosphate-buffered saline (BSA/PBS) solution. Atlantis T3 C18 column was used in HPLC (Waters, USA) for the separation of Trp metabolites with binary gradient solvent consisting of solvent A mmol/l ammonium acetate) and solvent B (methanol). The flow rate was 0.25 ml/min, and the injection volume was 10 µL. fractions were separated with positive and negative mode. Mass spectrometry was equipped with an electrospray source. All seven metabolites and their internal standards were infused into mass spectrometry for scanning the spectrum and determining fragmentation. Selected parent/fragment pairs for each uremic solute were used for mass spectrometry detection. Tryptophan, kynurenine, kynurenic acid, and xanthurenic acid were detected in positive mode, while indoxyl sulfate, indoxyl acetate, and anthranilic acid were detected in negative mode. The Institutional Review Board of Boston University Medical Center, MA approved the protocol.

### Mouse tissue homogenization

During necropsy, tissue samples were placed in 600 μL of RNA later (MilliporeSigma: #R0901500ML) and preserved at −80°C. Prior to RNA extraction, 20–30 mg of tissue was measured and transferred into a 2 mL tube containing 600 μL of RLT buffer with 1% β– mercaptoethanol and a 5 mm stainless steel bead (Qiagen, Hilden, Germany: #69989). The tissue was then dissociated using a Qiagen TissueLyser II (Qiagen) with the following cycle: two minutes of dissociation at 1800 oscillations/min, one minute of rest, followed by another two minutes of dissociation at 1800 oscillations/min. The dissociated tissues were centrifuged at 13,000 rpm for 10 minutes at room temperature, and the clear supernatant was carefully transferred to a new 1.5 mL Eppendorf tube before RNA extraction using TRIzol.

### Gene expression by RT-PCR

Total RNA was extracted with TRIzol (Life Technologies, Carlsbad, CA) and transcribed into cDNA using the QuantiTect Reverse transcription kit (Qiagen), and transcript levels were quantified by TaqMan PCR assays with commercially available primer sets (Applied Biosystems), by the ^ΔΔ^Ct method, with glyceraldehyde-3-phosphate dehydrogenase (GAPDH) as a normalizing control ^32^.

### Cell surface TF activity

As previously described, a two-step FXa generation assay was used to measure the TF procoagulant activity in ECs ^17,18,32^. In Brief, HUVECs were seeded in a 96-well plate (1000/well), serum-starved for 16 h, then treated with 5% sera from COVID-19 patients for 24h with and without AHR (CH223191; 10 µM) and IDO-1 (INCB024360; 10 µM) inhibitors. A standard plot was generated by incubating human recombinant lipidated TF (Enzolifesciences, Cat# SE-537) ranging from 0-500 pM along with 5 nM of human factor VIIa (Enzyme Research Laboratories Cat# HF-VIIa) and 150 nM of factor X (Enzyme Research Laboratories Cat# HFX 1010) and CaCl2-5 mM for 30 min at 37 °C. A chromogenic substrate was added for factor Xa (Chromogenix Cat# S2765, 1mM-final concentration) and then incubated for 5 min. The reaction was stopped using 10 µL of 50% glacial acetic acid and read at 405 nm absorbance.

### AHR activity

AHR activity was measured using HUVEC-TERT2 cells stably expressing a Cignal Lenti Reporter xenobiotic response element tethered to luciferase reporter (XRE-luc) (Qiagen, CLS-9045L-8), as previously described ^21^. In brief, the cells were seeded at 1×10^3^ cells per well in 96-well plates. Cells were starved for 16 hours and exposed to with and without AHR inhibitor CH223191 (Selleckchem, USA) in 5 % sera for 24 h. Firefly luciferase activity was measured using PROMEGA’s Dual Reporter Luciferase Assay Kit (cat#: E1980). The Luciferase signal (relative luciferase activity unit) was normalized by total protein (Bradford protein assay kit; ThermoFisher, MA, USA).

### IDO activity assay

IDO activity was determined, as described previously,^33,34^ and calculated by dividing the detected concentration of Kyn in serum by that of Trp.

### Immunohistochemistry and immunofluorescence

Following necropsy, mouse tissue samples were fixed for 72 hours in 10% neutral buffered formalin. Tissues were then processed in a Tissue-Tek VIP-5 automated vacuum infiltration processor (Sakura Finetek USA, Torrance, CA, USA), and subsequently, paraffin embedded using a HistoCore Arcadia paraffin embedding station (Leica, Wetzlar, Germany). Immunohistochemistry (IHC) and immunofluorescence (IF) were performed as previously described ^21^. In brief, paraffin embedded (liver, lungs, kidney, and aorta) tissue sections (5μm) were deparaffinized and antigen retrieval was performed using 1X antigen unmasking solution (Vector laboratories, H3300). For IHC, we used mouse and rabbit Specific HRP/DAB IHC Detection Kit–Micro-polymer (ab236466) for the detection of specific antigens followed by tissues counterstained with hematoxylin and eosin. The following antibodies used for IHC: Kynurenine 3-monooxygenase (KMO) (Abcam, ab220922); Tryptophan 2,3-dioxygenase (TDO2) (Millipore, MABN1537); Kynureninase (KYNU) (NOVUS, NBP1-56545); Indoleamine 2,3-dioxygenase (IDO-1) (Millipore, MABF850). Dilution factors for these antibodies were provided in **supplemental Table 1.**

For IF, tissues were permeabilized with 0.3% Triton X-100 in 1X phosphate-buffered saline (PBS) for 10 min, blocked with 5% bovine serum albumin, and primary antibodies were probed overnight at 4°C followed by secondary antibodies from Thermo Fisher Scientific (Alex Fluor 488, 594) for 45 min and counterstained with DAPI. The following antibodies used for IF: These antibodies were CD31 (Abcam, ab28364); TF (Abcam, ab104513); AHR (Thermo Fisher Scientific, MA1-514). Dilution factors for these antibodies were provided in **supplemental table 1.**

All images were obtained and quantified using the NikonQ17 Eclipse TE2000 Inverted Microscope at the Boston University School of Medicine (BUSM) Imaging Core. For quantification, the signal was converted to grayscale, and the number and intensity of pixels were analyzed as integrated density by using Fiji software (version 2.9.0). The integrated density of all the images was normalized to its area.

### Statistical analysis

Statistical analysis was performed with Prism 8 software (GraphPad) by one-way analysis of variance (ANOVA) for multiple groups and the Bonferroni multiple-comparisons test or Student *t* test for 2 groups.

## Results

### Alterations in the kynurenine pathway in liver, kidneys and lungs of humanized ACE2 mice

Since TDO in the liver is the first enzyme in Kyn biogenesis, we examined the liver of K18-hACE2 mice infected with SARS-CoV-2 at 2dpi, 4dpi, 7dpi, and mock controls. The expression was quantified using integrated density (ImageJ), which combines the intensity and pixel of signal and normalizes to fixed area^21,35^. IHC of liver cross-sections (**Figure 2A^a-d^**) and quantification (**Figure 2A^e^**) showed that TDO levels increased at 2dpi (~3-fold) and peaked at 4dpi (~5-fold) and reverted at 7dpi. Next, we examined other Kyn catabolic enzymes in **the** kidneys and lungs of K18-hACE2 mice infected with SARS-CoV-2. IHC of kidney cross-sections showed IDO-1 expression in the renal tubules (**Figure 3A^a-d^**), and its expression increased ~3-fold in SARS-CoV-2-infected mice at 2dpi and remained elevated until 7dpi compared to controls (**Figure 3A^e^**). Subsequently, we tested downstream enzymes in Trp catabolism, such as Kynurenine-3-monooxygenase (KMO) and L-kynureninase (KYNU). Interestingly, both KMO (**Figure 3B^a-d^ and Figure 3B^e^**) and KYNU (**Figure 3C^a-d^ and Figure 3C^e^**) were downregulated in SARS-CoV-2 infected mice at 4dpi and 7dpi compared to control kidneys.

**Figure 2:**
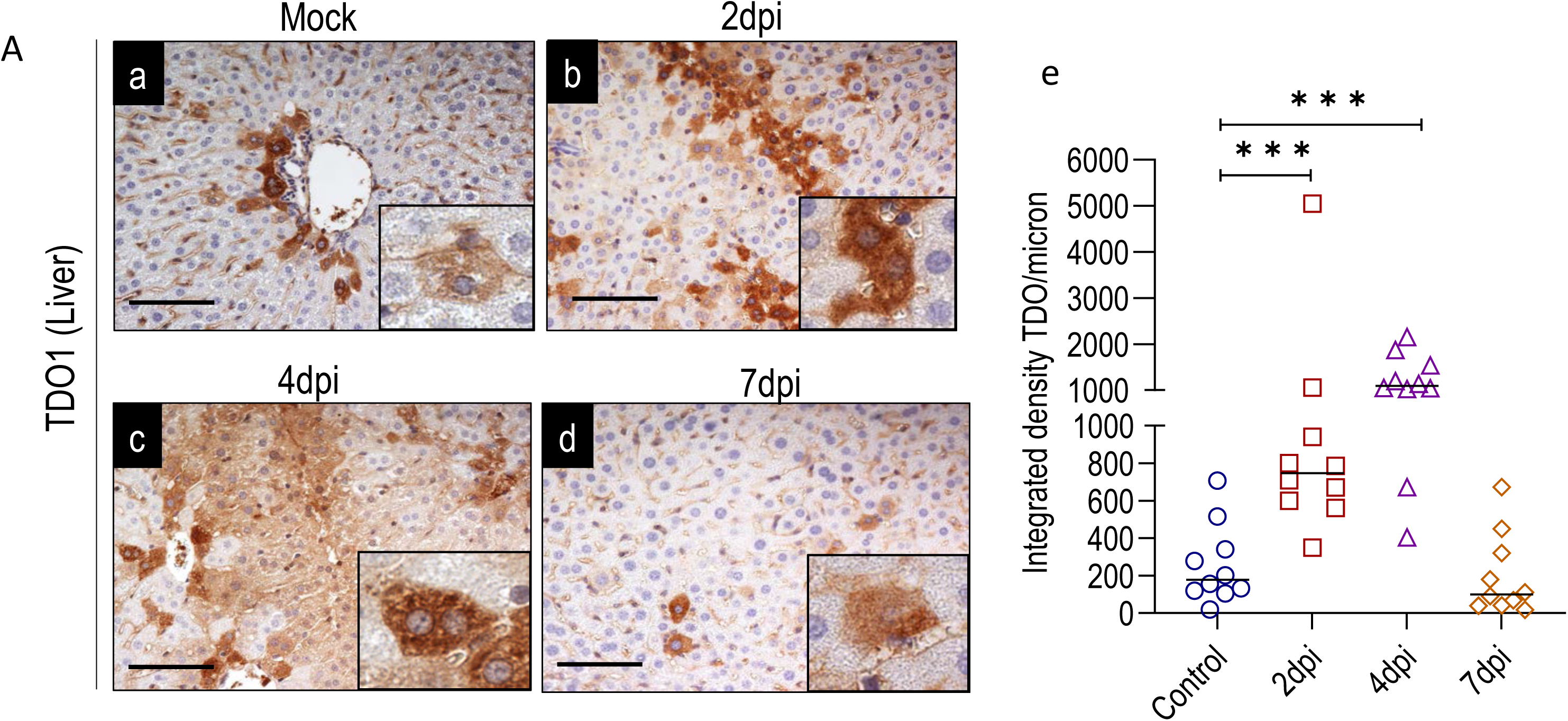
TDO regulation in liver of humanized transgenic ACE2 mice infected with SARS-CoV-2. Mice were infected with SARS-CoV-2 and paraffin-embedded liver sections were made at 2dpi, 4dpi, and 7dpi and screened for TDO-1 immunoreactivity (n=3 and imaged 3 different locations per mouse). (a) Represents mice exposed to normal saline that serves as a mock control; (b) 2dpi; (c) 4dpi, and (d) 7dpi. Scale bar = 100 micron; the inserts depict the zoomed-in images. (e) Data were quantified and plotted as integrated density per micron. Data were analyzed one-way ANOVA followed by multiple comparison test*. Significant ***P< .005*.

**Figure 3:**
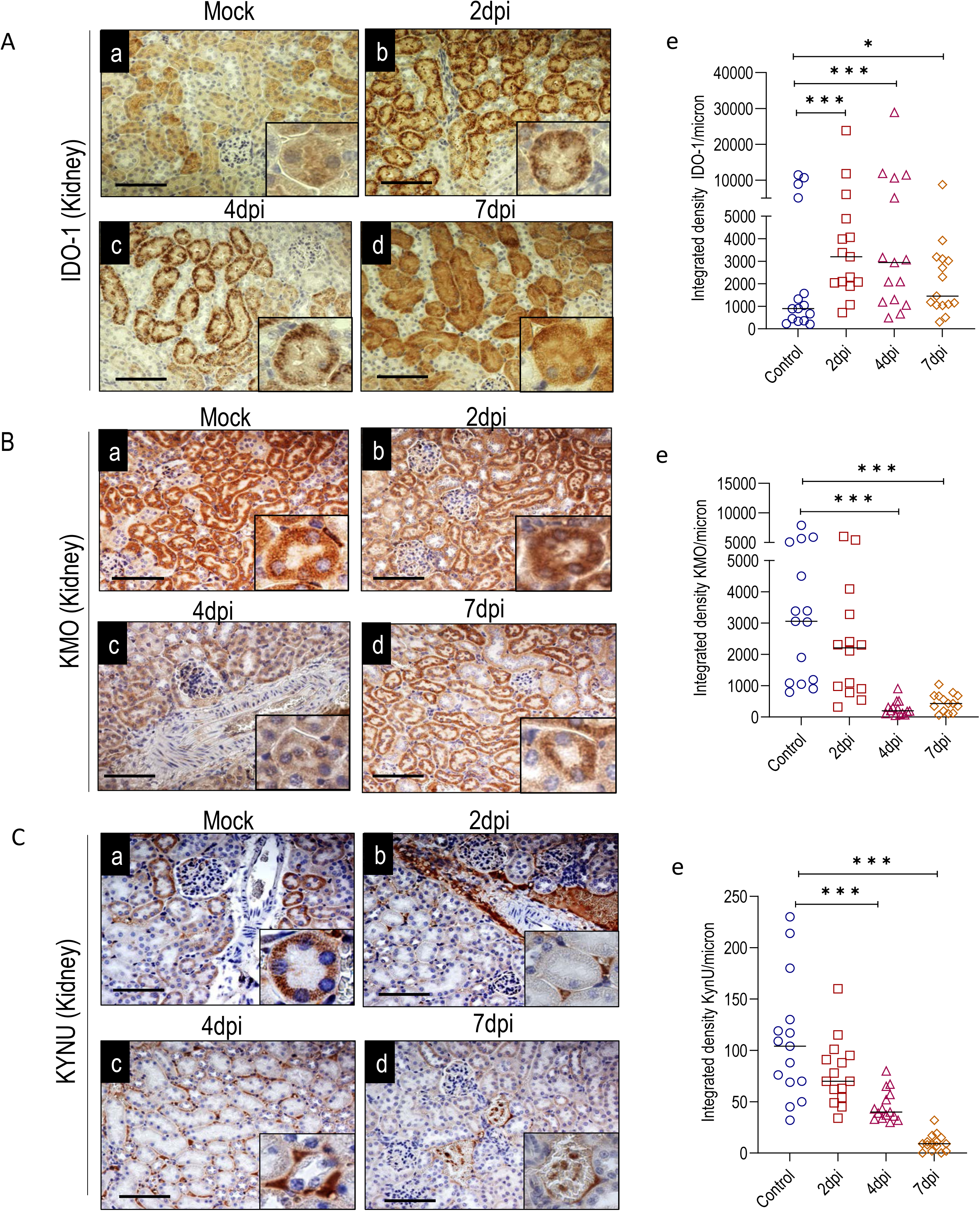
IDO-1, KMO, and KYNU expression in renal tubules of K18-hACE2 mice infected with SARS-CoV-2. Mice were infected with SARS-CoV-2 and paraffin-embedded kidney sections were made at 2dpi, 4dpi, and 7dpi and screened for (A) IDO-1, (B) KMO, and (C) KYNU immunoreactivity (n=3 and imaged 3 different locations per mouse) in renal tubules. (a) Represents the mice exposed to normal saline that serves as a mock control; (b) 2dpi; (c) 4dpi, and (d) 7dpi in all three figures (A, B, and C). Scale bar = 100 micron; the inserts depict the zoomed-in images. (e) Data were quantified and plotted as integrated density per micron of respective markers (A, B, and C). Data were analyzed one-way ANOVA followed by multiple comparison test. *Significant ***P< .005*.

A similar analysis was performed in the lungs of K18-hACE2 mice infected with SARS-CoV-2. IDO-1 expression in pulmonary epithelial cells was significantly upregulated at 2dpi (~ 2-fold) and peaked at 4dpi (~4-fold) and reversed back at 7dpi compared to mock controls (**Figure 4A^a-d^ and Figure 4A^e^**). KMO (**Figure 4B^a-d^ and Figure 4B^e^**) and KNYU (**Figure 4C^a-d^ and Figure 4C^e^**) expression significantly reduced in lungs at 2dpi, 4dpi and 7dpi compared to controls.

**Figure 4:**
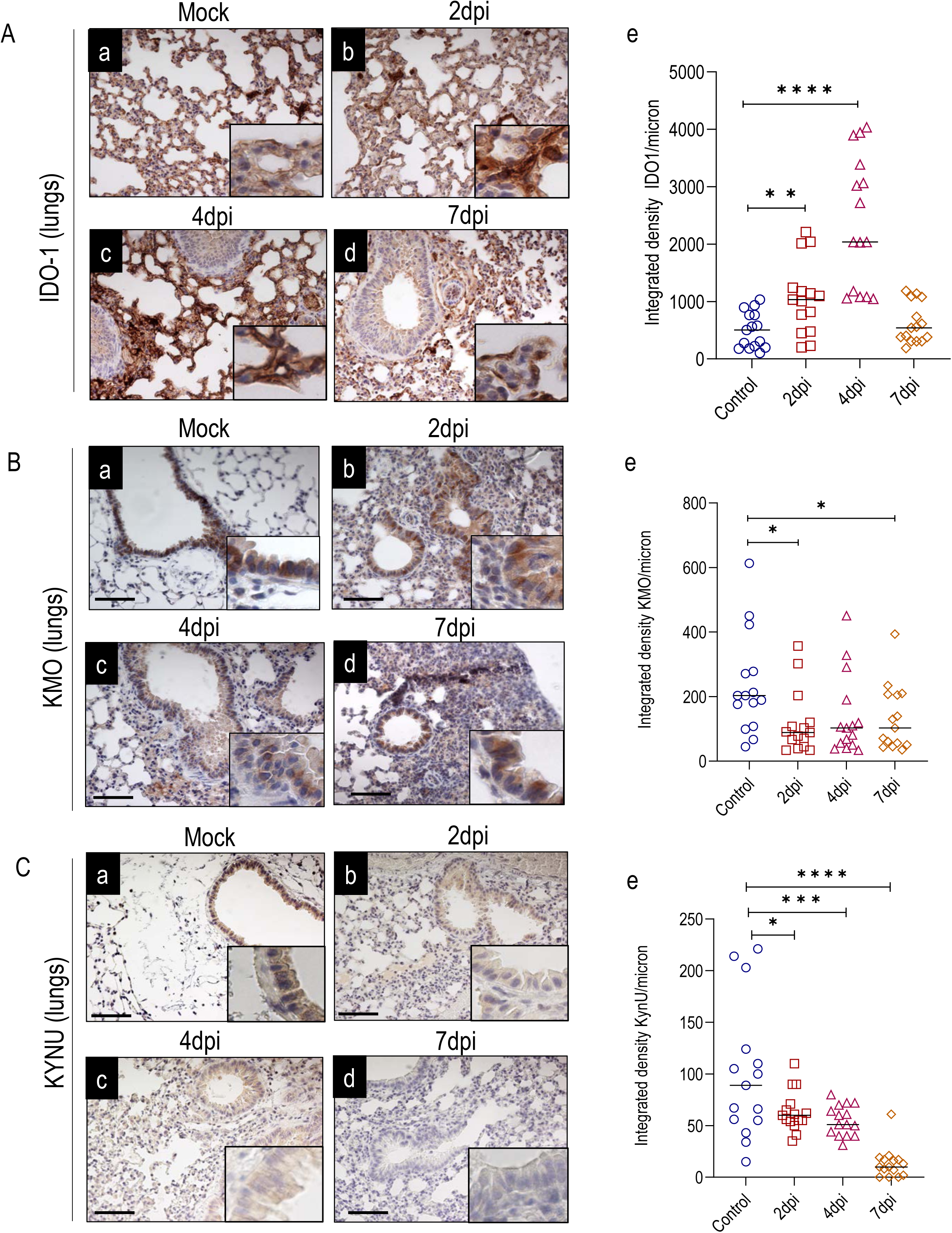
IDO-1, KMO, and KYNU expression in lungs of humanized transgenic ACE2 mice infected with SARS-CoV-2. Mice were infected with SARS-CoV-2 and paraffin-embedded lung sections were made at 2dpi, 4dpi, and 7dpi and screened for (A) IDO-1, (B) KMO, and (C) KYNU immunoreactivity (n=3 and imaged 3 different locations per mouse) in lungs. (a) Represents the mice exposed to normal saline that serves as a mock control; (b) 2dpi; (c) 4dpi, and (d) 7dpi in all three figures (A, B, and C). Scale bar = 100 micron; the inserts depict the zoomed-in images. (e) Data were quantified and plotted as integrated density per micron of respective markers (A, B, and C). Data were analyzed one-way ANOVA followed by multiple comparison test. *Significant ***P< .005*.

We further validated the expression of these enzymes using RT-PCR in liver, kidney, and lungs. RT-PCR data revealed that TDO mRNA expression in liver was upregulated after SARS-CoV-2 infection at 2dpi, 4dpi and significantly increased (~2-fold) at 7dpi compared to controls **(supplemental Figure 1A)**. IDO-1 expression in lungs, but not in kidney, was significantly upregulated at 2dpi and remained elevated at 4dpi and 7dpi compared to mock controls **(supplemental Figures 1B-C**). KMO and KYNU were downregulated in both kidney and lungs at 7dpi **(supplemental Figures 1D-G**). As changes in the expression of IDO-1 protein in kidney did not fully correspond to mRNA levels, these data suggest potential post-transcriptional regulation of IDO-1 enzyme in this tissue. These findings suggest the upregulation of TDO1 in hepatocytes and IDO-1 in kidney and lung ECs and the suppression of enzymes involved in KYN catabolism.

### Increase in Kynurenine biogenesis and activation of AHR signaling in SARS-CoV-2-infected mice

Our data showed upregulation of KYN biogenesis enzyme and downregulation of enzymes regulating KYN catabolism. This perturbation could result in an increase in KYN levels in the blood. Thus, we examined mouse sera samples at 4dpi (**Figure 5A**) and sera from COVID-19 patients (**Figure 6A**). Sera were analyzed using a pre-validated targeted metabolomics approach using Liquid Chromatography Mass Spectrometry (LC-MS/MS) (**Figure 5A**) ^18,36^. We observed a significant increase in KYN (~3 to 4-fold; **Figure 5B**), KYNA (~6-fold; **Figure 5C**), and AA (~2-fold; **Figure 5D**) in patients sera compared to non-infected controls. No significant changes in Trp (**Figure 5E**) and QA (**Figure 5F**) were noted.

**Figure 5:**
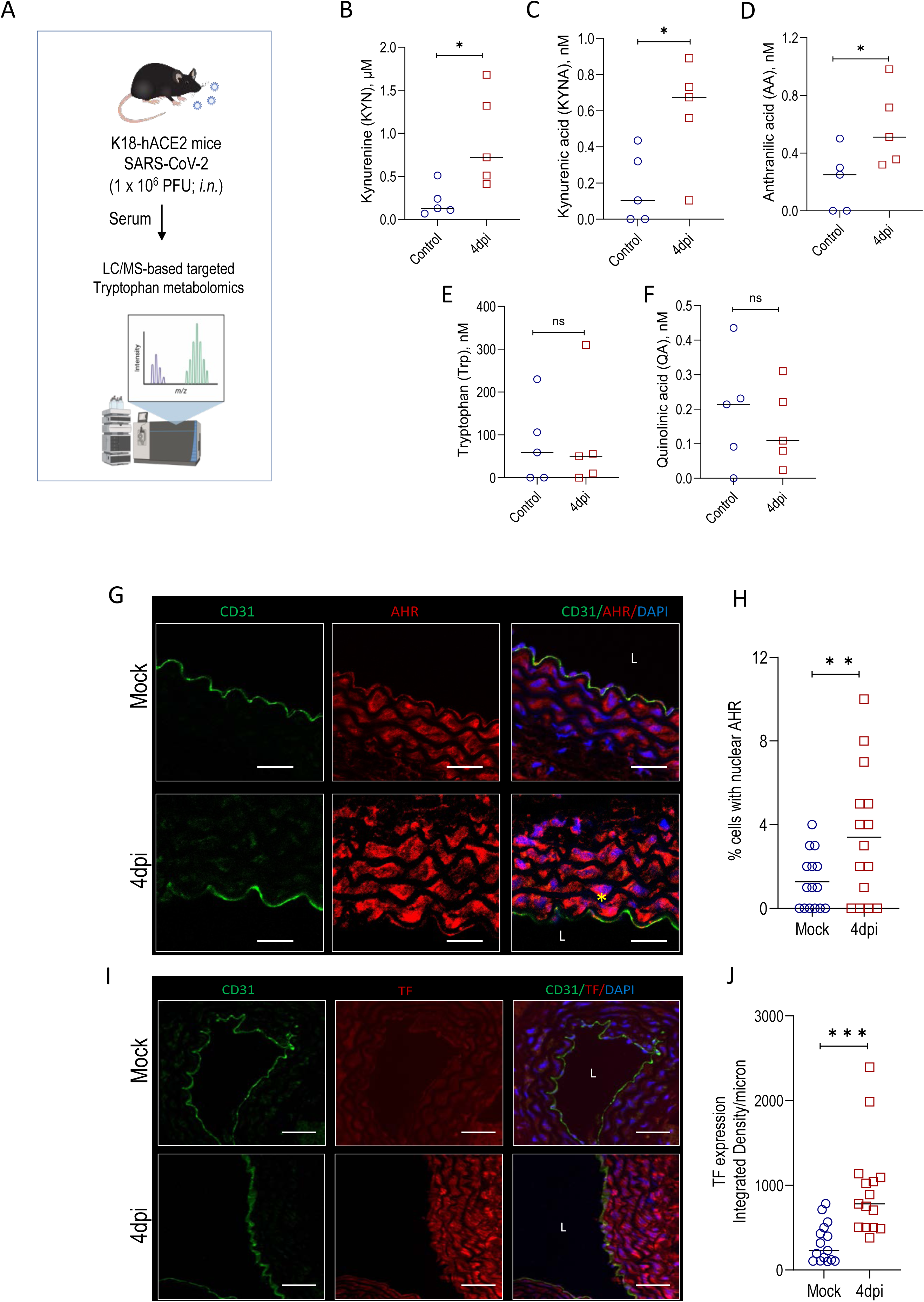
LC/MS-based targeted metabolomics in sera and AHR and TF expression in renal arteries from K18-hACE2mice (4dpi) Sera samples from K18-hACE2mice infected with SARS-CoV-2 were used for LC/MS-based metabolomic study (A). Separation was on Atlantis T3 C18 3µm 50 × 2.1 mm column Waters Corporation (MA, USA). A binary gradient consisting of solvent A (5 mmol/l ammonium acetate) and solvent B (methanol) and different gradients were applied (please see the method section for specific details about each metabolite). Mass spectrometry of each metabolite was examined using API 4000 triple quadrupole mass spectrometry equipped with electrospray and Agilent HPLC pump G1312B using plasma samples. All five metabolites and their internal standards were infused into mass spectrometry for scanning spectrum and determining fragmentation (B-F). All the metabolites were estimated from 4dpi sera and compared to mock controls (n=5 per group). Metabolites: Tryptophan (Trp), kynurenine (KYN), kynurenic mouse sera; *student t-test. *P < .05; n.s., not significant*. K18-hACE2 mice were infected with SARS-CoV-2 and paraffin-embedded kidney sections were made at 4dpi and stained for (G) AHR (I) TF. CD31 was used in both the panels (G and I) as an endothelial marker. Scale bar: panel A: 400X, Panel B: 200X). The yellow asterisk depicts the nuclear AHR expression, L: lumen. (H and J) Data were quantified and plotted as integrated density per micron. Data were analyzed by *student t-test*. *Significant *P< .05*.

**Figure 6:**
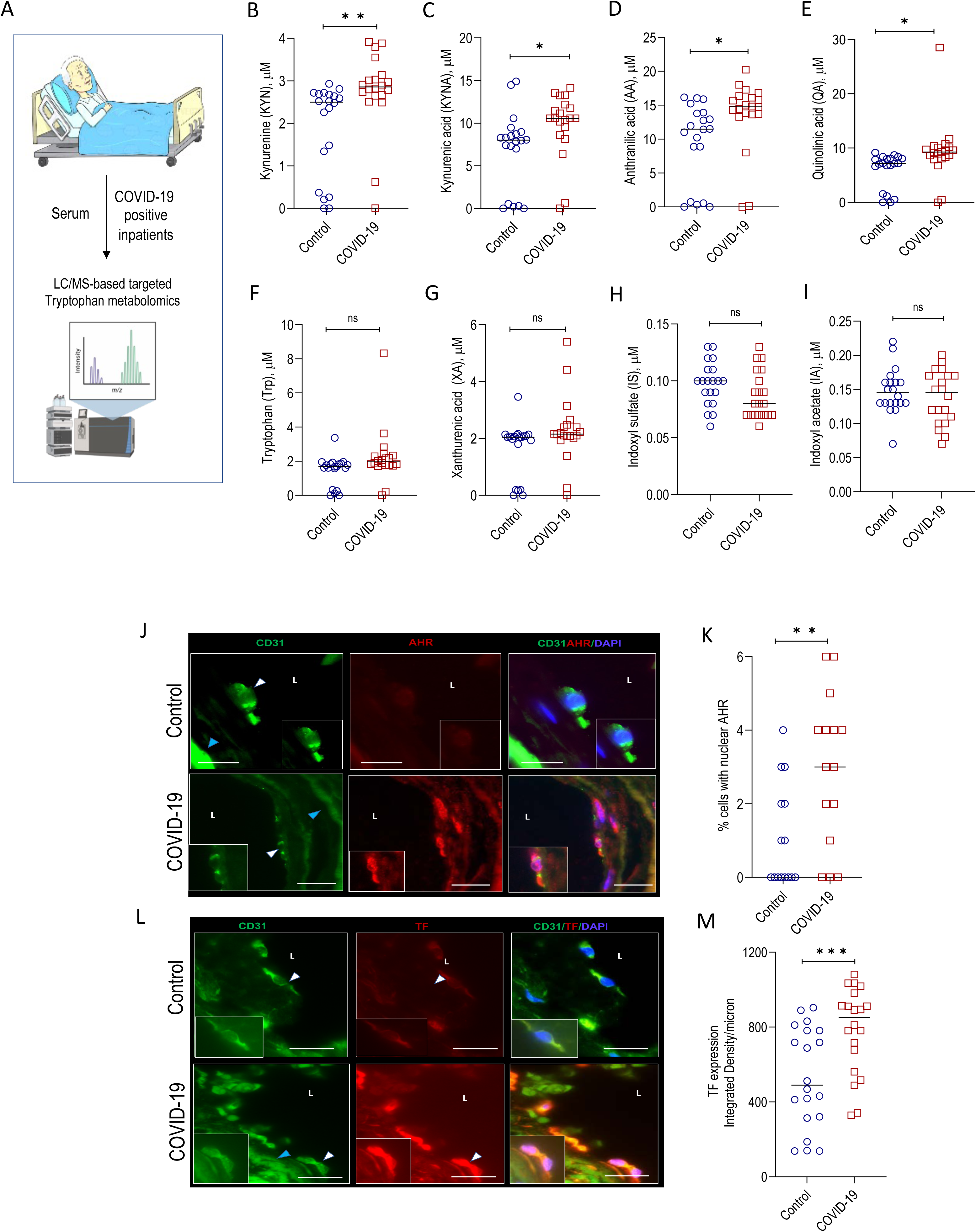
LC/MS-based targeted metabolomics in sera from COVID-19 patients and AHR and TF expression in renal arteries from human autopsy. Sera samples from COVID-19 patients and healthy controls were used for LC/MS-based metabolomic study (A). Separation was on Atlantis T3 C18 3µm 50 × 2.1 mm column Waters Corporation (MA, USA). A binary gradient consisting of solvent A (5 mmol/l ammonium acetate) and solvent B (methanol) and different gradients were applied (please see the method section for specific details about each metabolite). Mass spectrometry of each metabolite was examined using API 4000 triple quadrupole mass spectrometry equipped with electrospray and Agilent HPLC pump G1312B using plasma samples. All seven metabolites and their internal standards were infused into mass spectrometry for scanning spectrum and determining fragmentation (B-I). All the metabolites were estimated from COVID-19 patients’ sera and compared to matched controls (n=20). Metabolites: Tryptophan (Trp), kynurenine (KYN), kynurenic acid (KYNA), xanthurenic acid (XA), anthranilic acid (AA), quinolinic acid (QA), indoxyl sulfate (IS), indoxyl acetate (IA); n = 20; each dot represent individual patients’ sera; *student t-test*. \**P* < .05; \*\**P* <.01; *n.s.,* not significant. Paraffin-embedded kidney sections were made from COVID-19-infected human autopsy (64-YO) and stained for (J) AHR (L) TF. CD31 was used in both the panels (J and L) as an endothelial marker. Scale bar = 100 microns. Inserts show a zoomed-in image of the cells of interest. (K and M) Data were quantified and plotted as integrated density per micron in ECs. L= lumen; White and blue arrowheads show the ECs. Data were analyzed by *student t-test*. *Significant **P< .05; ***P< .005*.

Since the above metabolites (???) are known AHR ligands, we examined this hypothesis that AHR is activated in tissues of SARS-CoV-2-infected mice. Upregulation of nuclear AHR in cells and their target genes, such as CYP1A1 and CYP1B1, are considered telltale signs of AHR activation. Thus, we examined the mRNA expression of *Cyp1a1* and *Cyp1b1* by RT-PCR. *Cyp1b1* was significantly upregulated in kidney lysates after 4dpi (~3-fold) and remained elevated at 7dpi compared to mock controls **(supplemental Figure 2A).** Interestingly, we did see downregulation of *Cyp1ba1 expression (~*2 to 4*-*fold*) at all-time points* **(supplemental Figure 2B)**.

*Cyp1b1* was also significantly upregulated (~2-fold) in lung lysates after 2dpi and remain elevated at 4dpi and 7dpi compared to mock controls **(supplemental Figure 2B).** IF of kidney vasculature and quantification revealed a reasonable increase in AHR (~2-fold) in ECs and sub-ECs at 4dpi compared to mock controls (**Figure 5G-H**). Similarly, AHR expression was upregulated (~4-fold) in lung vasculature at 4dpi compared to mock controls **(supplemental Figure 3A-B**).

TF is a potent activator of the coagulation pathway when it is expressed on the cell surface^37^. We previously reported that the AHR-TF axis is associated with thrombosis after vascular injury in humans^17^ and in mouse cancer models subjected to venous thrombogenicity^21^. Thus, we sought to investigate whether SARS-CoV-2 infection alters TF expression in the kidneys and lungs of SARS-CoV-2-infected mice. RT-PCR revealed that TF mRNA expression was upregulated in mice in both kidneys and lungs at all-time points **(supplemental figure 4A-B**). IF of kidney vasculature revealed a reasonable increase in TF (~4-fold) in ECs and sub-ECs at 4dpi compared to mock controls (**Figure 5I-J**). Similarly, IF of lung vasculature showed TF upregulation (~2-fold) in lungs at 4dpi compared to mock controls **(supplemental Figure 3C-D**). Overall, these findings suggested that increased expression of AHR and TF in the vasculature could contribute to a prothrombotic milieu in the luminal region of K18-hACE2 mice infected with SARS-CoV-2. This implied that the vascular beds were inflamed, which could regulate activation of the extrinsic coagulation pathway via the AHR-TF axis.

### Kynurenine-AHR-TF axis in patients infected with SARS-CoV-2

To enhance translational relevance, we assessed the above observations in 20 patients confirmed to have SARS-CoV-2 infection between 2020-2022. Another group of patients negative for COVID-19 admitted to the same hospital during the same period served as controls. **Table 1** compares their baseline characteristics (see further details under methods).

COVID-19 patients had baseline heart rate, systolic blood pressure, and oxygen saturation levels similar to those of control patients. Moreover, white blood cell counts, and platelet counts were similar. Interestingly, ferritin was decreased in COVID-19-positive individuals, although the data were incomplete. There was a considerable elevation of D-dimer in COVID-19-positive patients, along with an increase in fibrinogen levels. About 40% of COVID-19 patients experienced respiratory failure, 20% of patients died, and 27.5% of patients had renal failure. It is important to highlight that patients’ sera samples were taken at the time of admission, and their creatinine levels were normal. They had not experienced unusual symptoms up to that point.

Interestingly, by LC-MS/MS targeted proteomics, we observed a significant increase in Kyn (~2-fold; *p= .009*), kynurenic acid (~2-fold; *p= .003*), anthranilic acid (~1.5-fold; *p= .028*), and quinolinic acid (~0.5-fold; *p= .020*), in COVID-19 patients sera compared to controls (**Figure 6B-E)**. We did not find any change in tryptophan, xanthurenic acid, indoxyl sulfate and indoxyl acetate concentrations in sera of COVID-19 patients compared to controls (**Figure 6F-6I**). Overall, these findings indicated systemic effect on Kyn catabolism that could potentially affect the luminal side of the vasculature, primarily ECs.

### Elevated TF and AHR in ECs in Kidney autopsy from COVID-19 patients

We examined the kidney autopsies from five COVID-19 patients to validate the findings in clinical settings. Kidneys of the controls and COVID-19 autopsies were examined, and tissues were stained to detect AHR and TF. IF data revealed a basal expression of TF in controls and a significant increase in AHR (~ 3-fold; *p= .006*) (**Figure 6J-K**) and TF (~2-fold; *p= .002*) **Figure 6L-M**) expressions in kidney EC of COVID-19 autopsies compared to controls.

### COVID-19 sera-mediated TF activation in human ECs can be suppressed by AHR or/and IDO-1 inhibitors

Next, we posited that sera of COVID-19 patients with high Kyn levels can activate AHR signaling in EC, thereby upregulating TF-dependent prothrombotic activity. First, we examined AHR activity of pooled sera from COVID-19 patients in primary HUVECs. Sera samples from COVID-19 (n = 29) and control (n = 31) patients were combined respectively and added to cultured ECs that stably express the AHR response element coupled to luciferase reporter. Promega Dual Luciferase (PDL) assay (Promega, USA) was used to analyze the lysates after ECs were treated with 5% sera for 16 h (**Figure 7A**). ECs exposed to COVID-19 sera exhibited significantly high AHR activity (~1.5-fold; *p= .043*) compared to control group sera (**Figure 7B**). Similarly, there was a significant increase in surface procoagulant TF activity (~2-fold*; p= .0001*). IDO-1 activity in sera was measured by the ratio of Kyn/Trp (see methods), which was ~1.5-fold; *p= .045* with COVID-19 sera compared to control sera (**Figure 7C-D**). TF activity induced in response to COVID-19 sera samples significantly correlated with AHR activity (*p= .045*) and IDO-1 activity (*p= .008*). (**Figure 7E-F**). Importantly, TF activity in ECs induced by sera of COVID-19 patients strongly correlated with the Kyn levels (*R*^2^ *= .562, P= .0001*) (**Figure 7G**).

**Figure 7:**
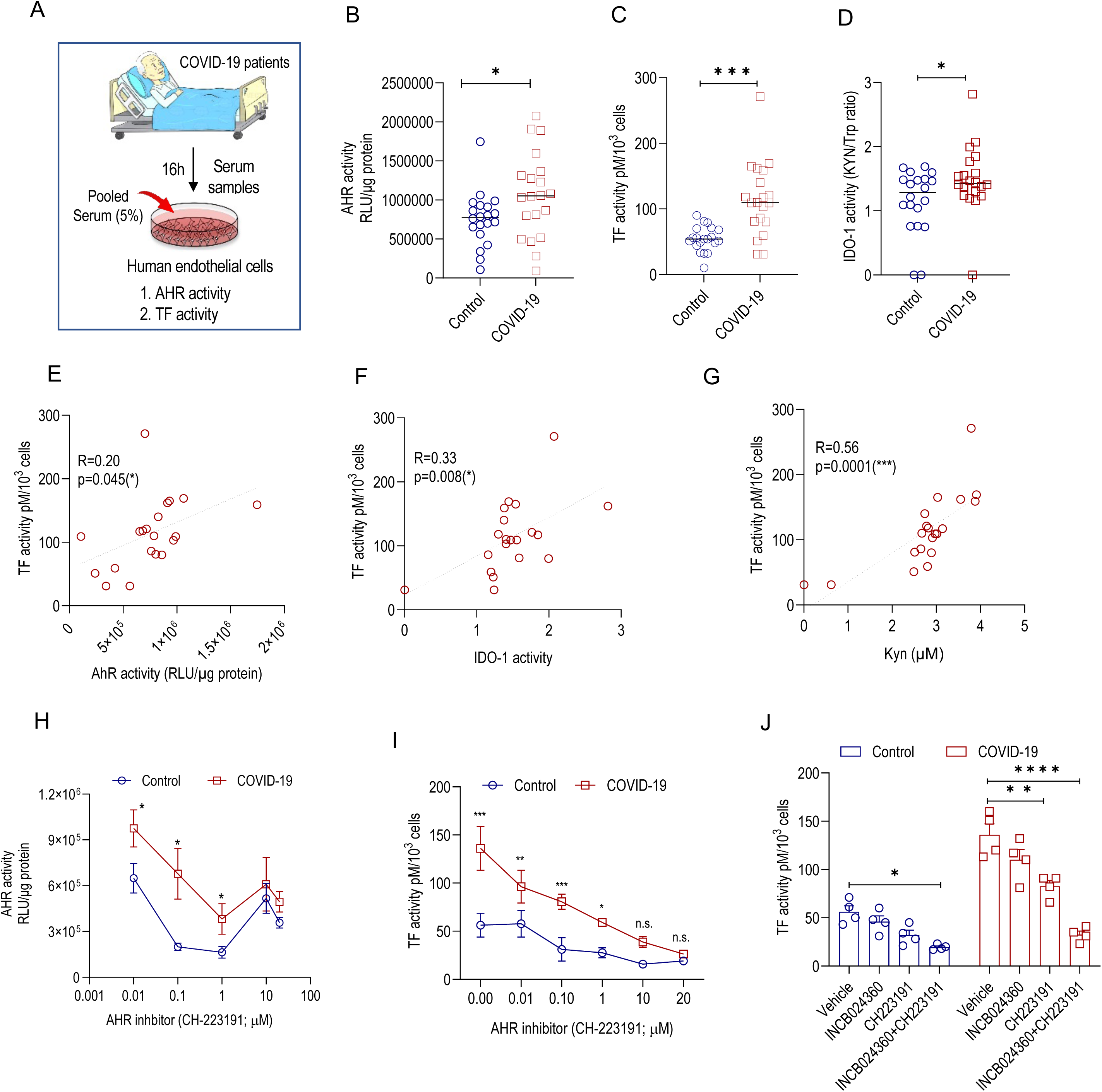
COVID-19 sera triggers AHR and TF activities in ECs. **(A)** HUVEC-TERT2 were exposed (16h) to COVID-19 patients’ pooled sera (5%) were obtained from BU biorepositories and measured AHR and TF activities. ECs that were genetically engineered stably expressing AHR response aliment tethered to a luciferase reporter. Cells were subjected to 5% sera obtained from control and patients with COVID-19. All the baseline characteristics of patients were shown in supplemental table 1. Cells were exposed to sera for 16h and AHR activity was performed using luciferase test. Data represent the individual samples with mean. *Significant **P< .01 by student t-test.* **(B)** AHR activity in control and COVID-19 patients sera exposed ECs. **(C)** TF activity in control and COVID-19 patients sera exposed ECs. **(D)** IDO-1 activity (sera levels of KYN/Trp). **(E)** Spearman’s rank correlation between TF activity and AHR activity. **(F)** Spearman’s rank correlation between TF activity and IDO-1 activity. **(G)** Spearman’s rank correlation between Kyn levels and TF activity. **(H)** AHR activity in control and COVID-19 patients subjected to the different concentrations (1nM to 100µM) of AHR inhibitor (CH223191) and ECs were pre-treated with control or COVID-19 sera along with AHR inhibitor (n=6). 2-way ANOVA for multiple groups with time scale observations followed by Bonferroni post-hoc test; *Significant, *P < .05; n.s., not significant*. Red circles represent WT; blue squares represent knock-out. **(I)** Pooled control and COVID-19 sera samples (5%) were exposed to ECs for 16h with AHR inhibitor (CH223191; 0 to 20µM) and measured cell-surface TF activity. Data were analyzed two-way ANOVA followed by multiple comparison test. *Significant ***P< .005*. Data were expressed as mean ± sem. n = 4; *significant *P < .05*; **(J)** Pooled control and COVID-19 sera samples (5%) were exposed to ECs for 16h with AHR inhibitor (CH223191; 10µM) and IDO-1 inhibitor (INCB024360; 10µM) and measured TF activity. Data were expressed as mean ± sem. n = 6. Data were analyzed two-way ANOVA followed by multiple comparison test. *Significant ns-non-significance, ***P< .001*.

As a pharmacological approach, we used COVID-19 and control group sera (5%) with increasing concentration of AHR inhibitor (CH223191) and examined AHR activity in ECs expressing AHR luciferase reporter. There was a significant dose-dependent reduction in AHR activity in both control and COVID-19 patients (**Figure 7H**).

To further study the AHR-TF axis, we treated human primary ECs with COVID-19 or control group sera (5%) with and without IDO-1 (INCB024360; 10µM) AHR (CH223191; 10µM) inhibitors and measured TF activity. At baseline, COVID-19 sera alone induced a ~3-fold higher TF activity in ECs compared to control sera (**Figure 7I**). AHR inhibitor significantly reduced endothelial TF activity upon both COVID-19 and control sera exposure (**Figure 7J**). However, we did not find notable changes in TF activity in ECs with IDO-1 inhibitor in the cell system (**Figure 8J**). It is possible that IDO-1 enzyme has no effect when a potent upstream regulator of TF, AHR has a dominant influence. In accordance, a co-inhibition approach (both AHR and IDO-1 inhibitors) resulted in ~6-fold reduction in endothelial TF activity (**Figure 7J**) in injected samples compared to controls.

**Figure 8:**
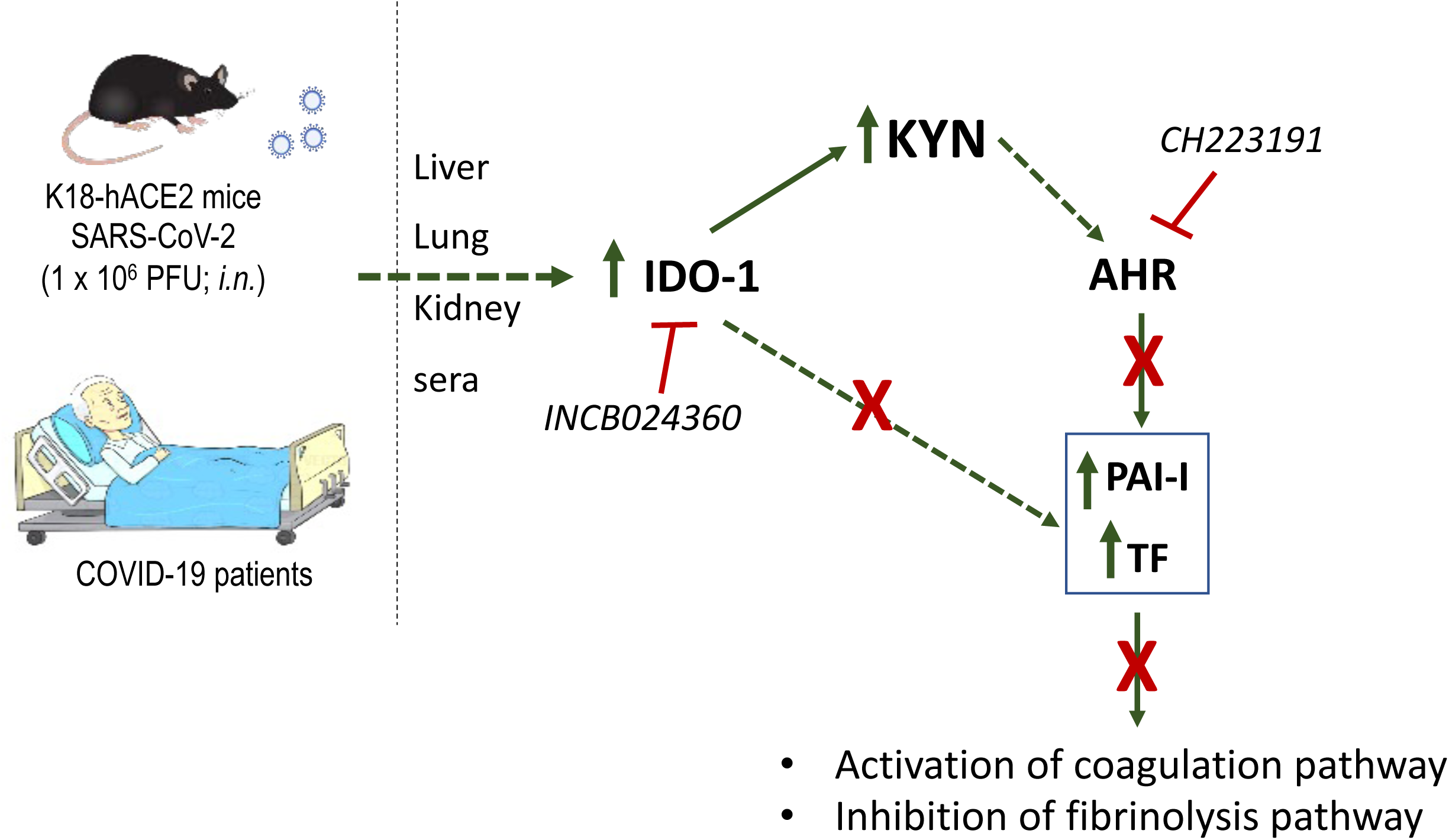
SARS-CoV-2-induced upregulation of IDO-1-Kyn-AHR axis induces a prothrombotic vasculature. Upregulated expression of IDO-1 in the vasculature of SARS-CoV-2-infected mice or humans, and resulting elevation of serum Kyn, an AHR activator, result in augmented levels of PAI-I and TF, the process of which is more effectively inhibited with both IDO-1 and AHR inhibitors; *green color arrow and signs indicates activation of Kyn pathway during SARS-CoV-2 infection in mice and human ECs; red color signs indicates inhibition of pathway by small molecule inhibitor (IDO-1 and AHR inhibitors)*.

Together, our results imply that cell-surface TF activity in ECs exposed to COVID-19 sera is mediated by AHR ligands in the sera, which were synergistically inhibited by IDO-1 and AHR inhibitors.

## Discussion

Our data showed that SARS-CoV-2 infection in mice and humans upregulated Trp catabolism due to differential regulation of key enzymes in Trp pathway in separate organs. Sera of patients with COVID-19 showed higher levels of Kyn, and a specific set of downstream catabolites. These results can be explained by the upregulation of TDO in liver, and IDO-1 enzyme in the lungs and kidneys of SARS-CoV-2 infected mice, supporting an increase in Kyn biogenesis in systemic (blood) and local milieu. This increase in Kyn levels is further accentuated by the downregulation of KUNU and KMO in the lungs and kidneys, suppressing the catabolism of Kyn. Our work also showed that sera samples from SARS-CoV-2-infected mice and humans activated AHR and TF activity in the ECs, which were suppressed by combined inhibition of AHR and IDO-1 (**Figure 8**). Overall, this study demonstrates a previously unrecognized reprogramming of Trp metabolism and activation of the AHR-TF axis in the ECs of lungs and kidneys in the context of SARS-CoV-2. This data also suggests a combinatorial effect of AHR and IDO-1 inhibitors as candidate-targeted antithrombotics in SARS-CoV-2 infected patients.

### Enzymes in Trp pathway in infections

Trp and its metabolites regulate different biological processes, particularly in infection and infection-related inflammation ^38–42^. Enhanced Trp catabolism is associated with infection-induced inflammation in *Chlamydia psittaci*, herpes simplex virus (HSV)-2, *Leishmania donovani*, and *Toxoplasma gondii* ^43^. However, these investigations did not compare the expression or regulation of different enzymes in Trp metabolic pathway. Our work uncovers alterations in key rate-limiting enzymes in Trp pathway such as IDO-1. *IDO-1* gene is regulated by interferon-α (IFNα), interferon γ (IFNγ), TNF-α, and prostaglandins ^44–47^. Recent work from our group showed that K18-hACE2 mice infected with SARS-CoV-2 had significant upregulation of plasma interferons (IFN-α/β/γ) and cytokines (TNF-α and IL-6) levels at 2dpi and 4dpi ^48^, which is likely to explain upregulation of *IDO-1* mRNA. Of note, KYNU and KMO expressions were downregulated without alterations in their respective mRNAs, suggesting a potential post-transcriptional regulation of these enzymes. Overall, our data suggest an interesting dichotomy in Kyn biogenesis and catabolizing enzymes and their differential regulation in SARS-CoV-2 warranting further investigation.

Changes in the enzymes revealed an interesting pattern over the post-infection period and may explain the chronology of thrombotic events in humans following SARS-CoV-2 infection. The baseline expression of TDO in the liver was low and was induced for 2-4 days post-infection, followed by suppression for seven days. In contrast, the expression of IDO-1 in renal tubules remained persistently elevated for seven days post-infection. The baseline expression KMO and KYNU was high in the kidneys and increased immediately post-infectious. However, their expressions were suppressed from post-infectious day four onwards. These results suggest that at baseline, Kyn metabolic homeostasis is biased towards its degradation. However, there is a change in Kyn metabolism following SARS-CoV-2 infection. In the immediate period following infection, enzymes in Kyn biogenesis and catabolism are increased simultaneously suggesting a potential net neutral or mild increase in levels of Kyn. However, the elevation of Kyn levels is sustained beyond four days post-infection by its increased biogenesis and reduced degradation. The increase in the local levels of Kyn in specific organs is likely to trigger the downstream signaling processes, such as the AHR-TF axis in the ECs of these organs, augmenting the risk of thrombosis in specific organs following SARS-CoV-2. The analysis of Wuhan COVID-19 cases showed systemic complications contributed by microvascular thrombosis, such as acute kidney injury or respiratory failure in the second week of SARS-CoV-2 infection^49^. Cases from New York also revealed events of arterial and venous thromboembolism between 8-14 days of initiation of symptoms of COVID-19^50^. Overall, our data suggest a dynamic nature of Trp metabolism in different organ systems, which may dictate the thrombotic risk in different organs.

### AHR-TF axis

Our data advances the observations of other reports. A recent report demonstrated that AHR is activated by SARS-CoV-2 infection to promote viral replication ^51^. Another study of human autopsies (66 patients with moderate to severe disease state) from COVID-19 patients demonstrated that upregulation of pulmonary endothelial TF and downregulation of thrombomodulin, a negative regulator of thrombosis, which correlated with elevated procoagulant markers in blood ^52^. Our results now show AHR as target of elevated Kyn in SARS-CoV-2 infection, with consequent upregulation of TF expression in ECs and induction of the extrinsic coagulation cascade.

### Translational significance

The translational relevance of our findings is underscored by the fact that prothrombotic metabolites, such as Kyn, elicit potential targetable pathways such as AHR. Also, metabolite generation can be pharmacologically manipulated. Both these approaches were explored to manipulate TF activity assay on ECs. Because a AHR inhibitor suppressed TF activity more than IDO-1 inhibitor, and both these inhibitors, when combined, had a higher TF suppressive effect, it raises a possibility of AHR activation by ligands other than Kyn or its catabolites, i.e., independent of IDO-1-mediated ligands. It is also possible that IDO1 inhibitor did not reduce Kyn to a threshold level that is needed to avoid AHR activation. The notion of combinatorial antithrombotic stands to reason, as prothrombotic propensity in COVID-19 patients requires a combination of therapies. Also, targeting such pathways, which do not interfere with physiological hemostasis mechanisms, is less likely to suffer from the major bleeding risk observed with Heparin^19,53^. These metabolites can be explored further as a potential biomarker guiding therapy with IDO-1 and AHR inhibitors as proposed in other human studies^34^

### Extended studies and conclusions

This study employed pharmacological manipulation, which might affect several cell types in context of whole-body physiology. Conditional knockout mice are required to establish a cell-type-specific contribution of different enzymes to SARS-CoV-2-induced thrombosis. Secondly, our study examined a set of Kyn degradation enzymes in two organs. Further investigation of other organs and the entire enzyme machinery of tryptophan is needed to comprehensively understand perturbations of Kyn metabolism. Similar to SARS-CoV-2, enveloped single-stranded RNA viruses, such as Ebola, Marburg, Lassa, South American, yellow, Crimea-Congo, and Rift Valley, cause prothrombotic complications^54^. Further studies are required to investigate the generalizability of the role of tryptophan metabolites and the AHR-TF pathway in other infections. SARS-CoV-2 has undergone several waves of mutations. It is important to compare the alterations in Trp metabolism with mutant SARS-CoV-2 and wild-type SARS-CoV-2 as the thrombotic risk is still observed with the mutant SARS-CoV-2^55^.

In conclusion, SARS-CoV2’s prothrombotic milieu can be explained by alterations in Trp metabolism and activation of AHR-TF axis in ECs of specific organs. In doing so, we demonstrate two anti-thrombotic pharmacological targets, AHR and IDO-1 in SARS-CoV-2, which supports further in vivo studies and pre-clinical testing as additional therapies for coagulation abnormalities in SARS-CoV-2 and possibly other viral related infections with thrombotic complications.

## Author contributions

V.C conceptualized the overall study. V.C., K.R., and S.S. designed the experimental study and offered conceptual insights. M.A.N., S.S., S.L., performed experiments and immunofluorescence studies. F.D. and D.K. performed mouse infections and provided tissue samples. M.H.K and H.K compiled COVID-19 patients’ details. V.J., and M.E. quantified immunohistochemistry slides. N.L. and S.W. performed a metabolomics study. G.Z., and E.B performed autopsy studies. C.A., M.B., and J.F. contributed conceptually to experimental studies. S.S., and V.C. interpreted data. S.S. wrote the manuscript. S.S., V.C., and K.R. reviewed and revised the manuscript. All authors read and approved the manuscript.

## Acknowledgment

We thank Prof Nigel Mackman (UNC Chapel Hill) for his assistance with the TF activity assay.

## Supplemental Figures

**Supplemental Figure 1:**
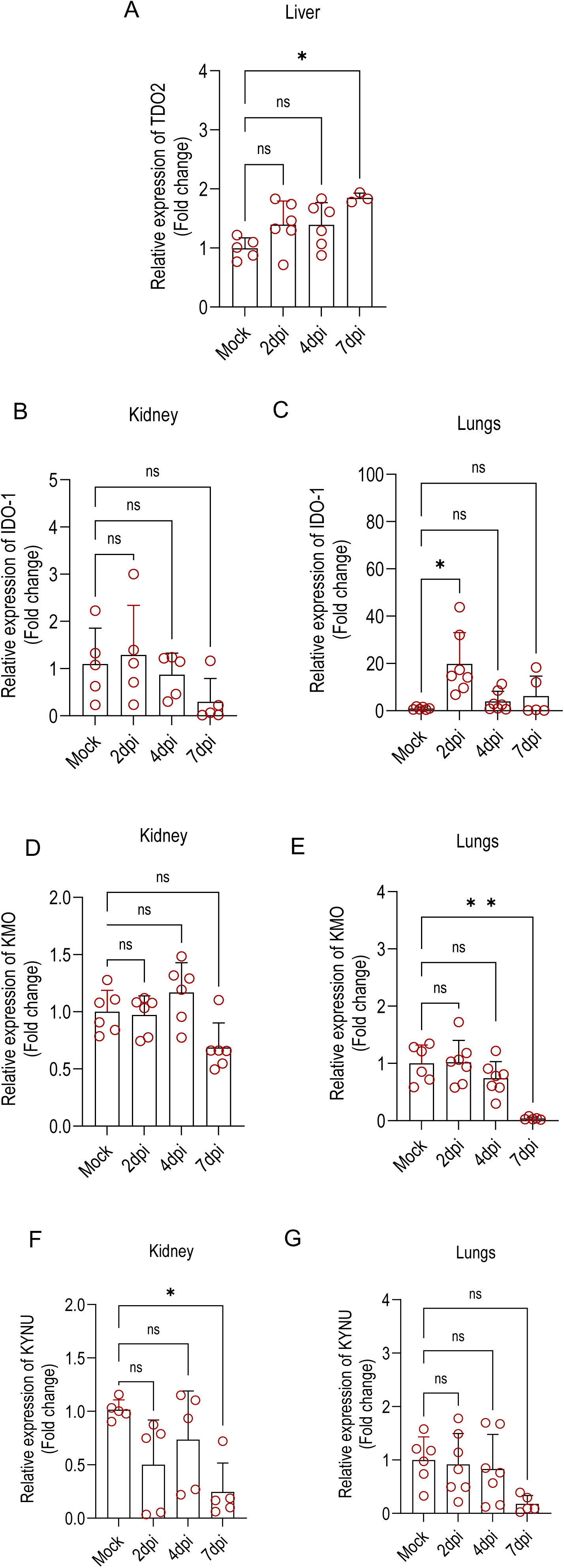
Gene expression in liver, kidney, and lungs from K18-hACE2mice infected with SARS-CoV-2. K18-hACE2 mice were infected with SARS-CoV-2 and RNA was isolated from liver, kidney, and lungs for gene expression by RT-PCR. (A) TDO2 expression in liver (B) IDO-1 expression in kidney (C) IDO-1 expression in lungs (D) KMO expression in Kidney (E) KMO expression in lungs (F) KYNU expression in kidney (G) KYNU expression in lungs. Data were analyzed one-way ANOVA followed by multiple comparison test. Data were expressed as mean ± SD; n=5-6 mice per group; *Significant: ns-non-significance, *P< .05, **P< .01*.

**Supplemental Figure 2:**
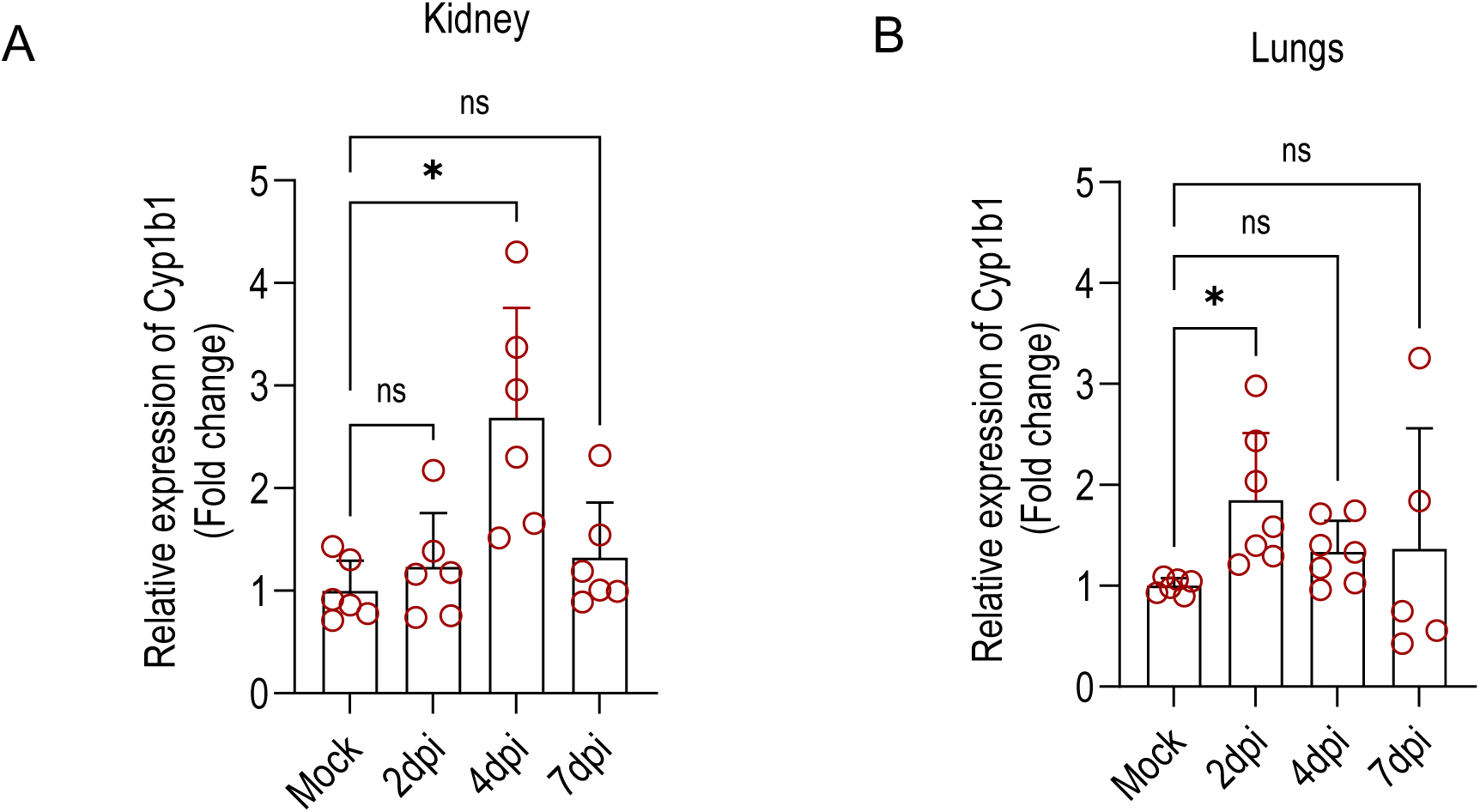
Cyp1b1 expression in kidney and lungs from K18-hACE2 mice infected with SARS-CoV-2. K18-hACE2 mice were infected with SARS-CoV-2 and RNA was isolated from kidney and lungs for cyp1b1 expression by RT-PCR. (A) Cyp1b1 expression in kidney (B) Cyp1b1 expression in lungs. Data were analyzed one-way ANOVA followed by multiple comparison test. Data were expressed as mean ± SD; n=5-6 mice per group; *Significant: ns-non-significance, *P< .05*.

**Supplemental Figure 3:**
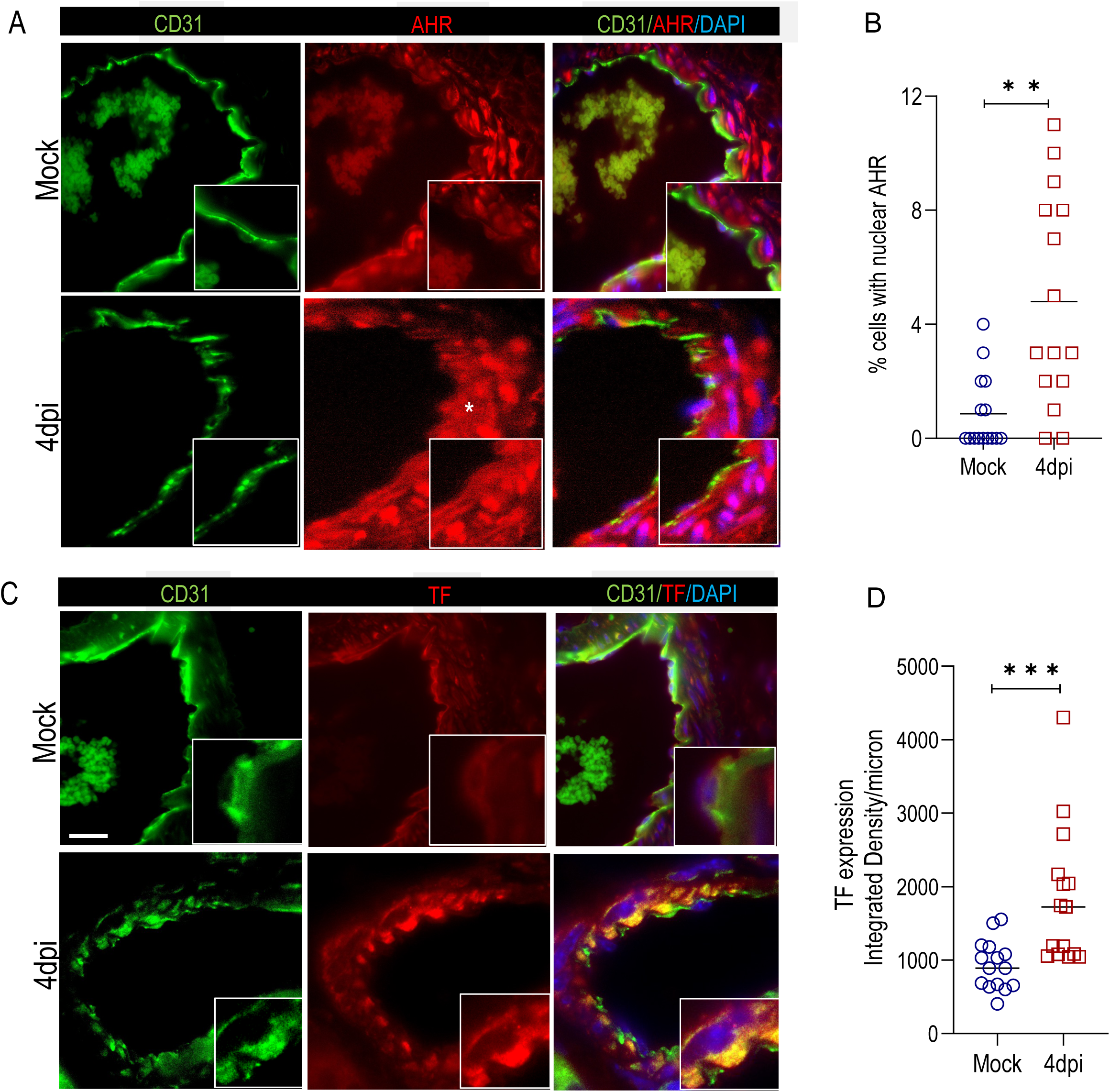
AHR and TF expression in lung vasculature from K18-hACE2 mice infected with SARS-CoV-2. K18-hACE2 mice were infected with SARS-CoV-2 and paraffin-embedded lung sections were made at 4dpi and stained for (A) AHR (C) TF. CD31 was used in both the panels (A and C) as an endothelial marker. Scale bar = 100 microns. The yellow asterisk depicts the nuclear AHR expression, L: lumen. (B and D) Data were quantified and plotted as integrated density per micron. Data were analyzed by *student t-test. Significant **P< .01; ***P< .001*.

**Supplemental Figure 4:**
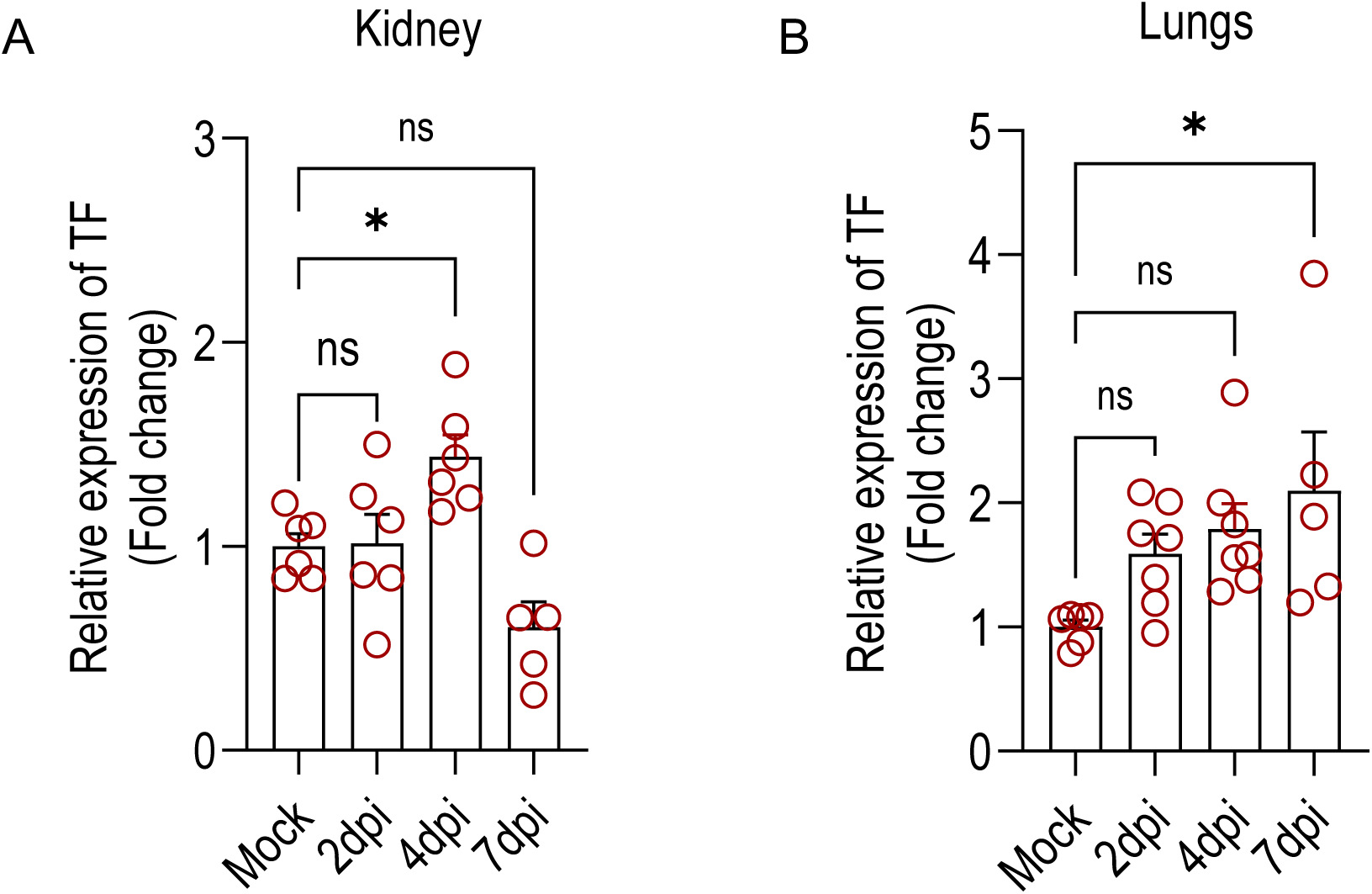
TF expression in kidney and lungs from K18-hACE2 mice infected with SARS-CoV-2. K18-hACE2 mice were infected with SARS-CoV-2 and RNA was isolated from kidney and lungs for TF expression by RT-PCR. (A) Cyp1b1 expression in kidney (B) Cyp1b1 expression in lungs. Data were analyzed one-way ANOVA followed by multiple comparison test. Data were expressed as mean ± SD; n=5-6 mice per group; *Significant: ns-non-significance, *P< .05*.

---

**Supplemental Table 1:** Antibodies used for Immunohistochemistry and immunofluorescence.

| Antibody | Vendor | Catalog number | Dilution |
| --- | --- | --- | --- |
| kynurenine 3-monooxygenase | Abcam | (ab220922) | IHC- 1:50 |
| Tissue Factor (TF) | Abcam | (ab104513) | IF- 1:100 |
| CD31 | Abcam | (ab28364) | IF- 1:50 |
| IDO (Clone 1F8.2) | Millipore | MAB10009 | IF- 1:10 |
| IDO-1 (Clone 4B7) | Millipore | MABF850 | IF- 1:10; IHC- 1:50 |
| Kynureninase | Novus | NBP1-56545 | IHC- 1:100 |
| TDO2 (clone7A6.1) | Millipore | MABN1537 | IHC- 1:50 |
| AHR | Thermo Fisher Scientific | MA1-514 | IF- 1:50 |

## References

1. Subramaniam S, Kothari H, Bosmann M. Tissue factor in COVID-19-associated coagulopathy. Thromb Res. 2022;220:35–47.

2. Kamel MH, Yin W, Zavaro C, Francis JM, Chitalia VC. Hyperthrombotic Milieu in COVID-19 Patients. Cells. 2020;9(11).

3. Mokhtari T, Hassani F, Ghaffari N, Ebrahimi B, Yarahmadi A, Hassanzadeh G. COVID-19 and multiorgan failure: A narrative review on potential mechanisms. J Mol Histol. 2020;51(6):613–628.

4. Vargas ESCF, Nguyen R, Willson M, et al. Intravital imaging of 3 different microvascular beds in SARS-CoV-2-infected mice. Blood Adv. 2023.

5. Qin Z, Liu F, Blair R, et al. Endothelial cell infection and dysfunction, immune activation in severe COVID-19. Theranostics. 2021;11(16):8076–8091.

6. Pellegrini D, Kawakami R, Guagliumi G, et al. Microthrombi as a Major Cause of Cardiac Injury in COVID-19: A Pathologic Study. Circulation. 2021;143(10):1031–1042.

7. Parra-Medina R, Herrera S, Mejia J. Systematic Review of Microthrombi in COVID-19 Autopsies. Acta Haematol. 2021;144(5):476–483.

8. Khismatullin RR, Ponomareva AA, Nagaswami C, et al. Pathology of lung-specific thrombosis and inflammation in COVID-19. J Thromb Haemost. 2021;19(12):3062–3072.

9. Ackermann M, Verleden SE, Kuehnel M, et al. Pulmonary Vascular Endothelialitis, Thrombosis, and Angiogenesis in Covid-19. N Engl J Med. 2020;383(2):120–128.

10. Mackman N, Grover SP, Antoniak S. Tissue factor expression, extracellular vesicles, and thrombosis after infection with the respiratory viruses influenza A virus and coronavirus. J Thromb Haemost. 2021;19(11):2652–2658.

11. Subramaniam S, Ruf W, Bosmann M. Advocacy of targeting protease-activated receptors in severe coronavirus disease 2019. Br J Pharmacol. 2022;179(10):2086–2099.

12. Conway EM, Mackman N, Warren RQ, et al. Understanding COVID-19-associated coagulopathy. Nat Rev Immunol. 2022;22(10):639–649.

13. Lin J, Yan H, Chen H, et al. COVID-19 and coagulation dysfunction in adults: A systematic review and meta-analysis. J Med Virol. 2021;93(2):934–944.

14. Wang L, He WB, Yu XM, Hu DL, Jiang H. Prolonged prothrombin time at admission predicts poor clinical outcome in COVID-19 patients. World J Clin Cases. 2020;8(19):4370–4379.

15. Subramaniam S, Reinhardt C, Kulkarni PP, Spiezia L. Editorial: COVID-19 and thrombo-inflammatory responses. Frontiers in Cardiovascular Medicine. 2023;10.

16. Chitalia VC, Shivanna S, Martorell J, et al. Uremic serum and solutes increase post-vascular interventional thrombotic risk through altered stability of smooth muscle cell tissue factor. Circulation. 2013;127(3):365–376.

17. Shivanna S, Kolandaivelu K, Shashar M, et al. The Aryl Hydrocarbon Receptor is a Critical Regulator of Tissue Factor Stability and an Antithrombotic Target in Uremia. J Am Soc Nephrol. 2016;27(1):189–201.

18. Kolachalama VB, Shashar M, Alousi F, et al. Uremic Solute-Aryl Hydrocarbon Receptor-Tissue Factor Axis Associates with Thrombosis after Vascular Injury in Humans. J Am Soc Nephrol. 2018;29(3):1063–1072.

19. Shashar M, Belghasem ME, Matsuura S, et al. Targeting STUB1-tissue factor axis normalizes hyperthrombotic uremic phenotype without increasing bleeding risk. Sci Transl Med. 2017;9(417):1–11.

20. Shashar M, Francis J, Chitalia V. Thrombosis in the uremic milieu--emerging role of “thrombolome”. Semin Dial. 2015;28(2):198–205.

21. Belghasem M, Roth D, Richards S, et al. Metabolites in a mouse cancer model enhance venous thrombogenicity through the aryl hydrocarbon receptor-tissue factor axis. Blood. 2019;134(26):2399–2413.

22. Savitz J. The kynurenine pathway: a finger in every pie. Mol Psychiatry. 2020;25(1):131–147.

23. Murakami Y, Saito K. Species and cell types difference in tryptophan metabolism. Int J Tryptophan Res. 2013;6(Suppl 1):47–54.

24. Capece L, Arrar M, Roitberg AE, Yeh SR, Marti MA, Estrin DA. Substrate stereo-specificity in tryptophan dioxygenase and indoleamine 2,3-dioxygenase. Proteins. 2010;78(14):2961–2972.

25. Dai X, Zhu BT. Indoleamine 2,3-dioxygenase tissue distribution and cellular localization in mice: implications for its biological functions. J Histochem Cytochem. 2010;58(1):17–28.

26. Cuffy MC, Silverio AM, Qin L, et al. Induction of indoleamine 2,3-dioxygenase in vascular smooth muscle cells by interferon-gamma contributes to medial immunoprivilege. J Immunol. 2007;179(8):5246–5254.

27. Beutelspacher SC, Tan PH, McClure MO, Larkin DF, Lechler RI, George AJ. Expression of indoleamine 2,3-dioxygenase (IDO) by endothelial cells: implications for the control of alloresponses. Am J Transplant. 2006;6(6):1320–1330.

28. Thomas T, Stefanoni D, Reisz JA, et al. COVID-19 infection alters kynurenine and fatty acid metabolism, correlating with IL-6 levels and renal status. JCI Insight. 2020;5(14).

29. Carossino M, Kenney D, O’Connell AK, et al. Fatal Neurodissemination and SARS-CoV-2 Tropism in K18-hACE2 Mice Is Only Partially Dependent on hACE2 Expression. Viruses. 2022;14(3).

30. Kenney DJ, O’Connell AK, Turcinovic J, et al. Humanized mice reveal a macrophage-enriched gene signature defining human lung tissue protection during SARS-CoV-2 infection. Cell Rep. 2022;39(3):110714.

31. Zhang A, Rijal K, Ng SK, Ravid K, Chitalia V. A mass spectrometric method for quantification of tryptophan-derived uremic solutes in human serum. J Biol Methods. 2017;4(3).

32. Subramaniam S, Ogoti Y, Hernandez I, et al. A thrombin-PAR1/2 feedback loop amplifies thromboinflammatory endothelial responses to the viral RNA analogue poly(I:C). Blood Adv. 2021;5(13):2760–2774.

33. Suzuki Y, Suda T, Asada K, et al. Serum indoleamine 2,3-dioxygenase activity predicts prognosis of pulmonary tuberculosis. Clin Vaccine Immunol. 2012;19(3):436–442.

34. Walker JA, Richards S, Whelan SA, et al. Indoleamine 2,3-dioxygenase-1, a Novel Therapeutic Target for Post-Vascular Injury Thrombosis in CKD. J Am Soc Nephrol. 2021;32(11):2834–2850.

35. Arinze NV, Yin W, Lotfollahzadeh S, et al. Tryptophan metabolites suppress Wnt pathway and promote adverse limb events in CKD patients. J Clin Invest. 2021.

36. Zhang A, Rijal K, Ng SK, Ravid K, Chitalia V. A mass spectrometric method for quantification of tryptophan-derived uremic solutes in human serum. J Biol Methods. 2017;4(3):1–8.

37. Mackman N. Role of tissue factor in hemostasis, thrombosis, and vascular development. Arterioscler Thromb Vasc Biol. 2004;24(6):1015–1022.

38. Mehraj V, Routy JP. Tryptophan Catabolism in Chronic Viral Infections: Handling Uninvited Guests. Int J Tryptophan Res. 2015;8:41–48.

39. Kavoussi L, Mikkelson D, Clayman R. Re: Rectal perforation as a complication of urethral instrumentation: 2 case reports. J Urol. 1989;142(5):1333–1334.

40. Ye Z, Yue L, Shi J, Shao M, Wu T. Role of IDO and TDO in Cancers and Related Diseases and the Therapeutic Implications. J Cancer. 2019;10(12):2771–2782.

41. Pantouris G, Mowat CG. Antitumour agents as inhibitors of tryptophan 2,3-dioxygenase. Biochem Biophys Res Commun. 2014;443(1):28–31.

42. Curti A, Trabanelli S, Salvestrini V, Baccarani M, Lemoli RM. The role of indoleamine 2,3-dioxygenase in the induction of immune tolerance: focus on hematology. Blood. 2009;113(11):2394–2401.

43. Schmidt SV, Schultze JL. New Insights into IDO Biology in Bacterial and Viral Infections. Front Immunol. 2014;5:384.

44. Werner ER, Bitterlich G, Fuchs D, et al. Human macrophages degrade tryptophan upon induction by interferon-gamma. Life Sci. 1987;41(3):273–280.

45. von Bergwelt-Baildon MS, Popov A, Saric T, et al. CD25 and indoleamine 2,3-dioxygenase are up-regulated by prostaglandin E2 and expressed by tumor-associated dendritic cells in vivo: additional mechanisms of T-cell inhibition. Blood. 2006;108(1):228–237.

46. Carlin JM, Borden EC, Sondel PM, Byrne GI. Interferon-induced indoleamine 2,3-dioxygenase activity in human mononuclear phagocytes. J Leukoc Biol. 1989;45(1):29–34.

47. Popov A, Abdullah Z, Wickenhauser C, et al. Indoleamine 2,3-dioxygenase-expressing dendritic cells form suppurative granulomas following Listeria monocytogenes infection. J Clin Invest. 2006;116(12):3160–3170.

48. Subramaniam S, Hekman RM, Jayaraman A, et al. Platelet proteome analysis reveals an early hyperactive phenotype in SARS-CoV-2-infected humanized ACE2 mice. bioRxiv. 2021:2021.2008.2019.457020.

49. Zhou F, Yu T, Du R, et al. Clinical course and risk factors for mortality of adult inpatients with COVID-19 in Wuhan, China: a retrospective cohort study. Lancet. 2020;395(10229):1054–1062.

50. Griffin DO, Jensen A, Khan M, et al. Arterial thromboembolic complications in COVID-19 in low risk patients despite prophylaxis. Br J Haematol. 2020.

51. Giovannoni F, Li Z, Remes-Lenicov F, et al. AHR signaling is induced by infection with coronaviruses. Nat Commun. 2021;12(1):5148.

52. Francischetti IMB, Toomer K, Zhang Y, et al. Upregulation of pulmonary tissue factor, loss of thrombomodulin and immunothrombosis in SARS-CoV-2 infection. EClinicalMedicine. 2021;39:101069.

53. Paranjpe I, Fuster V, Lala A, et al. Association of Treatment Dose Anticoagulation With In-Hospital Survival Among Hospitalized Patients With COVID-19. J Am Coll Cardiol. 2020;76(1):122–124.

54. Iba T, Levy JH, Levi M. Viral-Induced Inflammatory Coagulation Disorders: Preparing for Another Epidemic. Thromb Haemost. 2022;122(1):8–19.

55. Oba S, Hosoya T, Amamiya M, et al. Arterial and Venous Thrombosis Complicated in COVID-19: A Retrospective Single Center Analysis in Japan. Front Cardiovasc Med. 2021;8:767074.

